# Structural distance at the tRNA synthetase active site interface predicts pathogenicity but is captured by AlphaMissense and EVE except among score-ambiguous variants

**DOI:** 10.64898/2026.05.22.727252

**Authors:** Kayla Liebeskind, Christopher Francklyn, Ramiro Barrantes-Reynolds

## Abstract

Variants of uncertain significance have accumulated as genomic sequencing has become more widespread, which complicates rare disease diagnosis and requires substantial resources for re-evaluation. Aminoacyl-tRNA synthetases (ARSs) are a protein family with extensive variant data and well-characterized disease associations, making them an ideal system for investigating the relationship between variant location and pathogenicity. Using structural distance measurements to the ARS-tRNA binding interface combined with existing pathogenicity predictors, AlphaMissense and EVE, we investigated whether explicit structural binding information could improve missense variant pathogenicity prediction. Pathogenic variants were found to cluster significantly closer to the tRNA-binding interface than benign variants (p = 0.0003). Incorporating explicit distance information into a Bayesian mixture model did not substantially improve predictive performance over AlphaMissense and EVE alone, suggesting that these models already implicitly capture relevant structural binding context. However, a clinically important subset of interface variants classified as ambiguous by both existing models identifies a specific gap where explicit structural distance information may provide added discriminative value, but the limited number of clinically validated variants currently available constrains the ability to fully evaluate this potential. Incorporating additional biologically relevant features not captured by existing models, such as protein stability or conformational dynamics, as well as refining structural distance calculations, may further improve classification of this subset. These findings highlight both the power and the limitations of existing pathogenicity predictors and suggest that structurally informed approaches targeting the binding interface represent a promising direction for improving classification of these ambiguous variants that have great clinical significance.

**Author Summary:** Advances in clinical genetic sequencing have caused increasing identification of genetic variants whose impact on human health is unknown. These “variants of uncertain significance” present a major challenge because their role in causing disease cannot yet be confirmed or ruled out. This study focuses on a specific family of essential enzymes called aminoacyl-tRNA synthetases, which play a critical role in the process of proteins translation. Mutations in these enzymes have been linked to a range of diseases. This project aims to provide a novel method for determining pathogenicity of variants specifically in aminoacyl-tRNA synthetases. We propose that physical proximity of a variant to the functional binding site of the enzyme is influential in determining pathogenicity. We find that this spatial relationship is a meaningful indicator of a variant’s potential to disrupt normal function.

## Introduction

Aminoacyl-tRNA Synthetases (ARSs) are responsible for accurately coupling amino acids with their cognate tRNAs carrying specific anticodon sequences, which is essential for accurate translation of genetic information into functional proteins. This two-step reaction involves activation of an amino acid, followed by transfer of the amino acid to the 3’ end of the tRNA (1). Each ARS is named after the product generated, and most of these enzymes have both cytoplasmic and mitochondrial isoforms (2), but this project focuses on the cytoplasmic variants.

ARSs can be divided into class I and class II. Class I synthetases (CARS, EARS, IARS, LARS, MARS, QARS, RARS, VARS, WARS, YARS) share a Rossmann fold active site and typically function as monomers, approaching the tRNA acceptor stem from the minor groove to aminoacylate the 2’-OH. In contrast, class II synthetases (AARS, DARS, FARS, GARS, HARS, KARS, NARS, PARS, SARS, TARS) have an antiparallel β-sheet active site, generally function as dimers, and aminoacylate the 3’-OH from the major groove (3, 4).

Aminoacyl-tRNA synthetases generally recognize the inner L-shaped face of their cognate tRNAs, typically favoring nucleotide determinants located either in the acceptor stem (including A76, which is important in for elongation on the ribosome) or the anticodon region (nucleotides 34-36), which base-pairs with the mRNA codon for translation (5).

In mammals, eight cytoplasmic ARSs (RARS, DARS, QARS, IARS, LARS, KARS, MARS, and EPRS) associate with three auxiliary proteins to form the multi-tRNA synthetase complex (MSC). While the functional rationale of the MSC has not been unequivocally determined, prior research suggests that one possibility is that it serves to enhance the efficiency of charged tRNA delivery to the ribosome (6).

Beyond their canonical role in translation, ARSs participate in non-translational functions including transcriptional regulation, rRNA synthesis, and cell signaling. They also harbor editing domains that hydrolyze mischarged tRNAs to maintain translational fidelity, and eukaryotic ARSs frequently contain additional domains absent in bacterial homologs (3).

Defective ARS function is associated with a range of human diseases, as accurate aminoacyl-tRNA synthesis is essential for accurate protein synthesis because mischarged tRNAs can lead to amino acid misincorporation during translation (1, 7). Some pathogenic mutations in ARSs have either poorly defined effects on aminoacylation or uncertain impacts on enzyme stability, raising the possibility that such mutations affect non-canonical functions (8).

ARS mutations cause disease through both recessive and dominant mechanisms. Recessive mutations in cytoplasmic ARSs typically produce neurological phenotypes including hypomyelination, microcephaly, seizures, sensorineural hearing loss, and developmental delay, generally through loss of enzymatic function (9). Dominant mutations have been identified in five ARSs (GARS, YARS, AARS, HARS, and WARS), linked to Charcot-Marie-Tooth (CMT) disease, a peripheral neuropathy characterized by progressive motor and sensory loss. Because these ARSs function as homodimers, a mutant subunit can pair with a wild-type subunit to form a non-functional heterodimer, reducing overall activity below the threshold required by peripheral neurons. Owing to the fact that some of these mutations negatively impact the ability of the resultant heterodimeric ARS to release aminoacylated tRNA product, they phenotypically present as negative gain-of-function effects (10).

The growing records of disease-associated ARS variants has been documented in public databases such as ClinVar. ClinVar is a freely accessible NCBI archive that documents relationships between human genetic variants and disease, classifying variants as pathogenic, likely pathogenic, uncertain significance, likely benign, or benign based on submissions from clinical laboratories and research groups (11). However, these classifications are not definitive, as a substantial proportion of variants previously classified as pathogenic are reclassified upon re-evaluation, highlighting the inherent uncertainty in curated variant databases (12). UniProt similarly classifies variants using the same five-tier system, drawing on ClinVar, OMIM, and published literature (13, 14).

Despite rapid growth in variant databases enabled by advances in genome sequencing, the majority of variants in disease-related genes remain classified as variants of uncertain significance (VUSs). As of April 2024, VUSs comprised 44.6% of all 2.8 million germline variant records in ClinVar (15). Resolving these classifications is clinically important, as uncharacterized variants complicate treatment decisions and can expose patients to unnecessary interventions and their associated psychological and financial burdens (16).

Several experimental approaches have been developed to interpret VUSs. *In vitro* aminoacylation assays have confirmed loss-of-function effects for pathogenic variants in at least 15 ARS genes, providing strong evidence that reduced aminoacylation activity drives recessive disease. However, these assays require individual variant characterization and purified recombinant protein, which limits scalability (17). Humanized yeast complementation assays have similarly been used to evaluate AARS variants associated with dominant neuropathy, but these assays may not faithfully recapitulate the impacts of AARS mutations on differentiated eukaryotic cell types, most notably including neurons (10). *In vivo* models in mice, zebrafish, and *Caenorhabditis elegans* have validated that ARS loss-of-function reproduces human disease phenotypes, but each model has its own limitations (18).

Multiplexed assays of variant effect (MAVEs) are a higher-throughput alternative, as they are able to simultaneously evaluate thousands of variants by comparing sequencing frequencies before and after functional selection (19). However, MAVEs require specialized equipment, have so far been applied to only a small fraction of genes, and can miss pathogenic variants if experimental conditions do not accurately replicate the relevant biological context (20).

Because of these limitations, computational tools have become a valuable complement to experimental approaches. They can generate pathogenicity predictions at a much larger scale and can help to identify high-priority variants for experimental follow-up.

AlphaFold 2 and AlphaFold 3 have transformed structural biology by enabling accurate prediction of protein structures and molecular complexes from sequence alone (21, 22). However, both produce static predictions that do not account for ligand interactions or environmental conditions, and predictions can deviate from experimental structures at both global and local scales (23). Despite these limitations, AlphaFold predictions are central to understanding how genetic variants may impact protein function.

AlphaMissense uses the AlphaFold 2 architecture to predict the likelihood of a missense variant in a protein to cause disease. It uses three complementary sources of information to generate pathogenicity predictions: evolutionary context, structural context from AlphaFold 2, and population allele frequency data from human and non-human primate populations. It classifies variants into three categories based on a continuous pathogenicity score: “likely benign” (0–0.34), “ambiguous” (0.34–0.564), and “likely pathogenic” (0.564–1) (24). The output of the model is a scalar pathogenicity score found to have 90% precision when compared to ClinVar (25).

EVE (Evolutionary model of Variant Effect) is another pathogenicity prediction model. It is an unsupervised deep generative model that predicts variant pathogenicity entirely from evolutionary sequence data, without using clinical labels or structural information. Instead of learning from known pathogenic and benign variants, EVE learns purely from the patterns of natural sequence variation across species, avoiding the sampling biases of curated clinical databases (26).

AlphaMissense has been found to generally outperform EVE in accurate pathogenicity predictions, but both tools are still limited and vary by the specific gene and biological context. There are many other existing algorithms like SIFT, PolyPhen2, CADD, and REVEL that attempt to classify pathogenicity of variants, but no single pathogenicity predictor has been found to perform best in all settings (27).

Current pathogenicity prediction models do not explicitly account for structural binding information. For ARSs, whose primary function is tRNA binding, proximity to the binding interface is a plausible determinant of pathogenic impact. We hypothesized that variants closer to the ARS-tRNA binding site are more likely to disrupt binding and reduce aminoacylation activity, and therefore more likely to cause disease.

To test this, we compiled available cytoplasmic ARS structures, performed structural alignments to tRNA structures, and calculated variant distances to key tRNA features. We found that pathogenicity is negatively correlated with the summed distance from a variant to the terminal nucleotide and anticodon midpoint of the tRNA. We incorporated this structural information alongside existing pathogenicity scores from AlphaMissense and EVE into a machine learning model aimed at improving variant classification in ARSs. This approach was evaluated against current models to determine whether structural binding information provides additional predictive value.

## Results

### Pathogenic Variants Cluster Near the ARS-tRNA Interface

When looking at the structures of ARSs with pathogenicity highlighted using AlphaMissense scores, pathogenic variants appear to cluster on the surface of the ARS that has either been shown or predicted to interact directly with the inside face of the tRNA. The pathogenicity scores of mutations located at a farther distance from the predicted tRNA binding interface are lower, suggesting that they are more likely to be benign (Fig 1).

**Fig 1.**
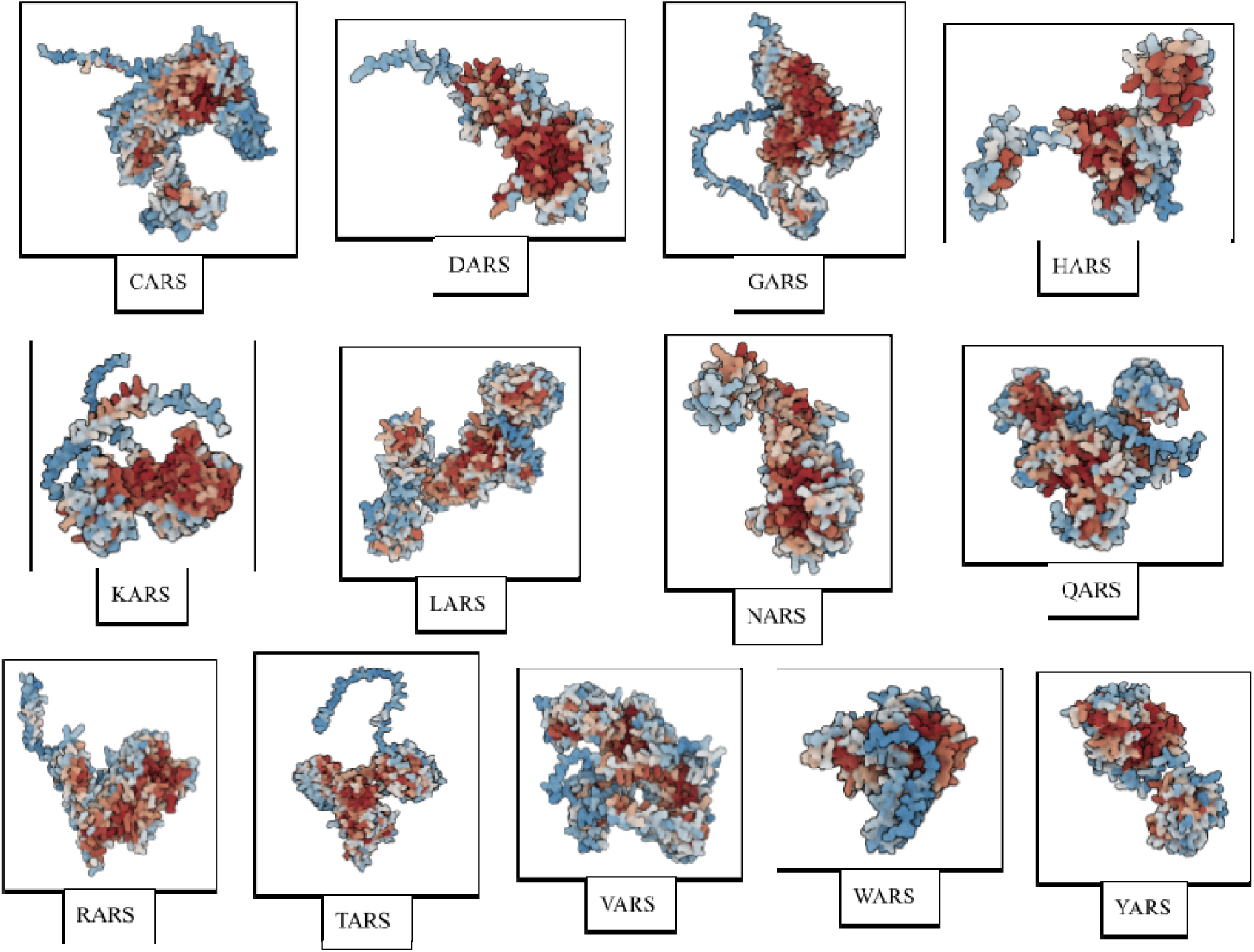
Structural overview with pathogenicity. Color shows average predicted pathogenicity in AlphaMissense for all possible substitutions at the position; blue residues are likely benign, gray residues are uncertain, and red residues are likely pathogenic. Structural models for each synthetase are derived from AlphaFold Protein Structure Database models (21, 28).

After aligning synthetase structures to their respective tRNAs, we calculated the minimum Euclidean distance from each residue in the ARS to key nucleotides in the tRNA. When compared to all other variants, pathogenic variants are found to cluster below a threshold of distance in three-dimensional space in relationship to the anticodon trinucleotide and the 3’ end of the tRNA. There are certainly other pathogenic variants outside of this threshold, but in general, pathogenic variants seem to be more likely to be found at distances below the threshold (Fig 2A). When comparing the sums of the distances to the two key nucleotides, pathogenic variants are significantly closer to these nucleotides than the benign variants (p = 0.0003) (Fig 2B).

**Fig 2.**
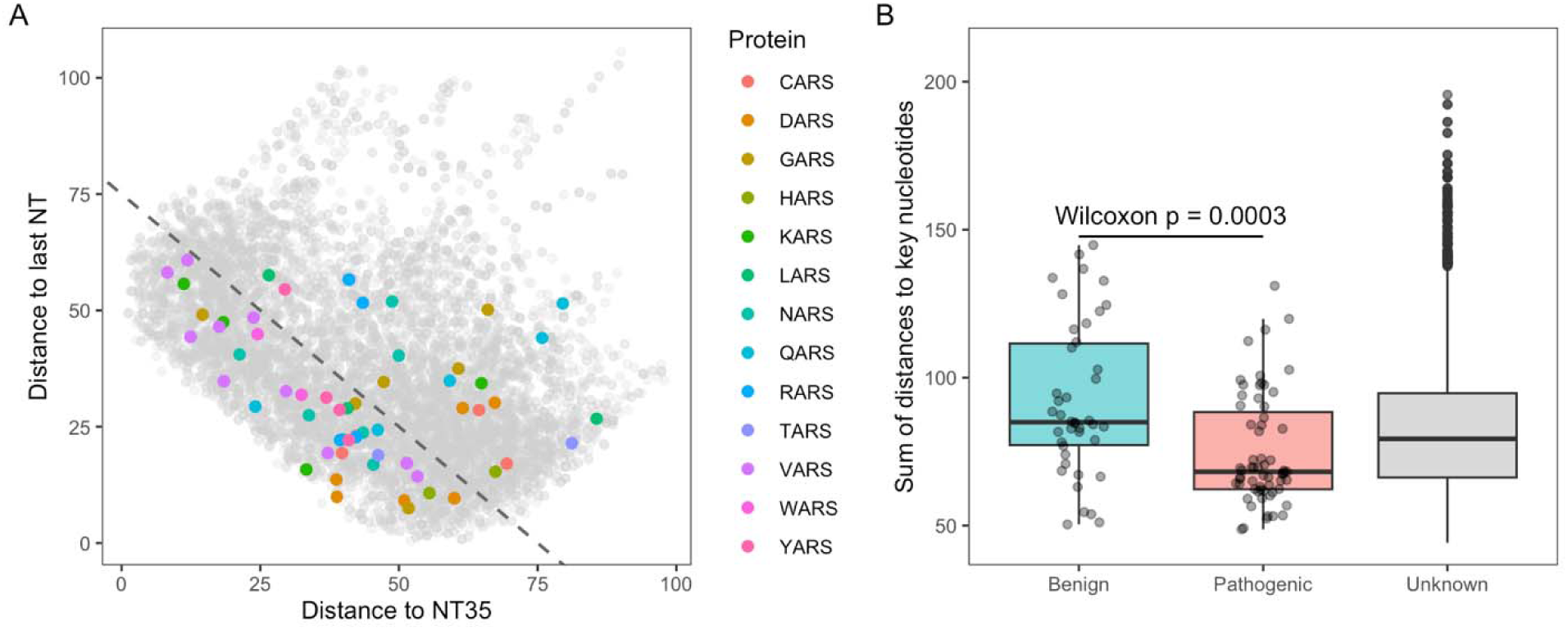
Mutation Euclidean distances (in angstroms) from variants to key nucleotides: tRNA anticodon (NT35) and nucleotide at the 3′ end of the acceptor stem (last NT, defined in density). (A) Distribution of pathogenic variants (colored by protein) and variants of unknown significance (gray) in proximity to the two key nucleotides. The dashed line represents a potential threshold for the interface region, corresponding to a sum of distances of 75 angstroms to the two key nucleotides: any residue below the diagonal would correspond to a position where d_1_+ d_2_ ≤ 75 angstroms. (B) Comparison of the sum of distances to the key nucleotides between pathogenic and benign variants. Variants of unknown pathogenicity are shown for reference.

We evaluated the quality of the sequence alignment using percent identity and percent similarity. HARS has the least accurate sequence alignment, with only 30.1% identity. Excluding HARS, alignment percent identity ranges from 75.2% to 91.7% (S3 Table).

Overall, there is extensive data about pathogenicity of mutations in ARSs (1, 17, 18, 29). There is much more information from predictive programs (EVE and AlphaMissense) than clinical data from UniProt (S1 Table). This reflects both the smaller number of identified variants in UniProt, which is limited to variants observed and characterized in the literature, as well as the exclusion of variants with conflicting or uncertain classifications from our dataset. Additionally, EVE analysis does not include five of the nineteen cytoplasmic aminoacyl-tRNA synthetases (including NARS, FARS, MARS, IARS, EPRS) of the 19 cytoplasmic synthetases. AlphaMissense has a higher average pathogenicity in this dataset than EVE (0.599 vs. 0.541, respectively). The median pathogenicity score for AlphaMissense (0.661) is also higher than that for EVE (0.552). Distance information is limited due to a limited number of proteins, but all mutations included in the dataset have calculated distance values to the two key nucleotides (the anticodon and 3’ end of the tRNA). The distance data lacks six synthetases: FARS, MARS, IARS, EPRS, SARS, and AARS (S1 Table).

The data are further divided by each individual synthetase studied. This includes only mutations with distance information. There are fewer EVE scores than AlphaMissense scores for each protein. Of the UniProt data, the majority of mutations are not classified as pathogenic or benign, they are either classified as having uncertain significance or have conflicting interpretations of pathogenicity; there are a total of 1035 mutations from UniProt, but only 63 are classified as pathogenic and 42 are classified as benign (S2 Table).

### Bayesian Mixture Model Combines Structural Distance with Existing Pathogenicity Predictors

Distances to key nucleotide 1 ranged from 1.275 to 97.925 angstroms, and distances to key nucleotide 2 ranged from 0.506 to 105.533 angstroms across all mutations in the dataset. Evaluating the data used for this model, a gamma distribution provided a good fit to the distribution of distances to both key nucleotides (Figs 3A and 3B, left for the PDF, and Figs 3A and 3B right for the CDF). Pathogenic variants in the synthetases cluster at slightly lower distances to both key nucleotides than benign variants (Figs 3A and 3B, right).

**Fig 3.**
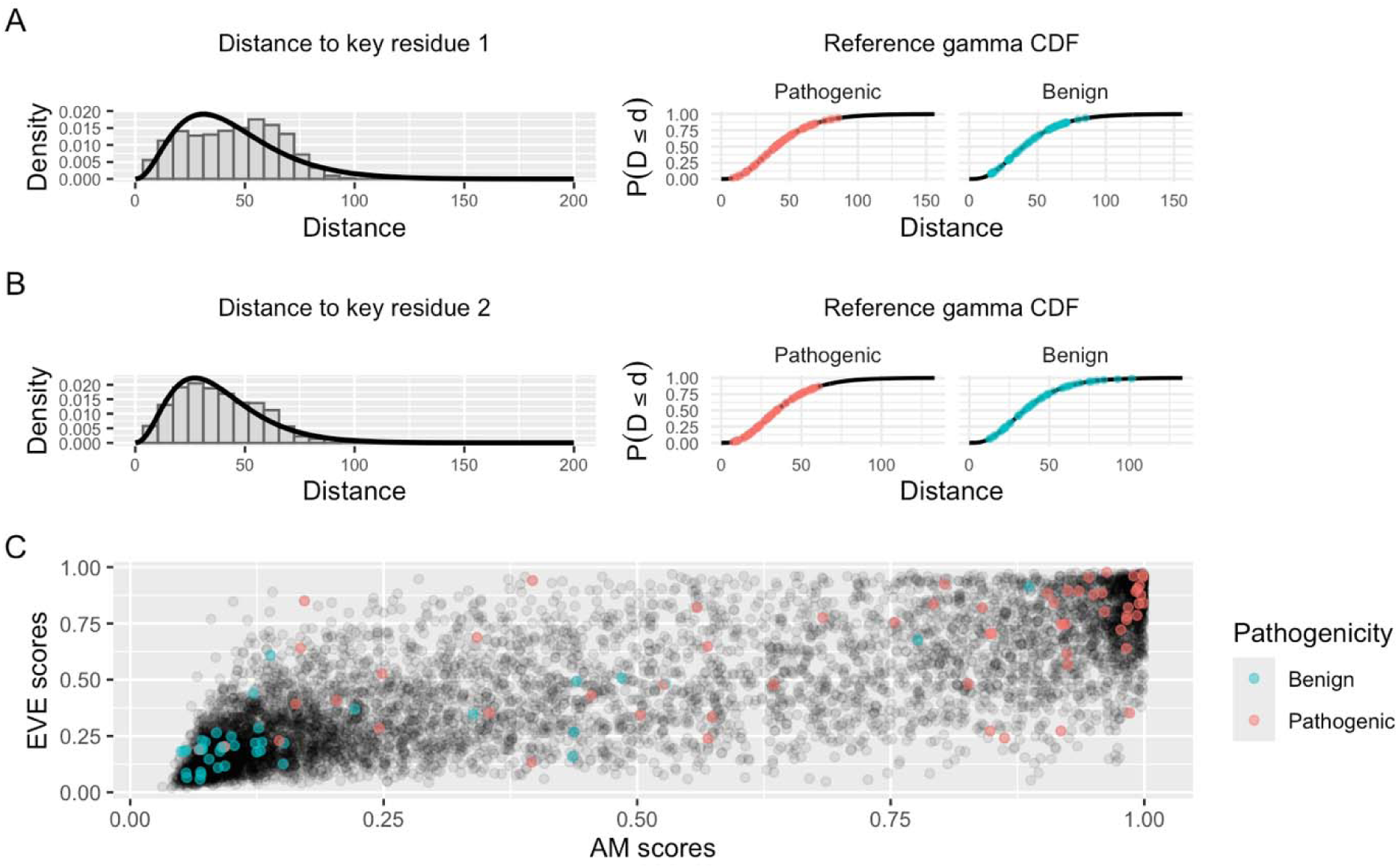
Evaluation of the data used for the new model incorporating structural distance information (in angstroms) into existing predictive models. (A) Top row. Left: Probability Density Function (PDF) of mutation distances to key nucleotide 1: tRNA anticodon (nucleotide 35). Right: Cumulative Distribution Function (CDF) of distances to nucleotide 1; pathogenic and benign mutations are highlighted separately. (B) Middle row. Left: PDF of mutation distances to key nucleotide 2: tRNA 3’ end (last nucleotide). Right: CDF of distances to nucleotide 2; pathogenic and benign mutations are highlighted separately. (C) Bottom row. Comparison of AM and EVE scores. Known pathogenicity variants are colored (blue for benign and red for pathogenic), and unknown variants are shown in gray.

A scatterplot comparing AlphaMissense and EVE scores reveals that in general, the two models agree on pathogenicity of variants, but there are several variants for which the methods disagree or are uncertain. Overall, AlphaMissense and EVE are highly correlated, especially for variants that are very likely pathogenic or benign, with spearman rank correlation

. They appear to disagree more on variants in the middle range of pathogenicity, between 0.30 and 0.70 (Fig 3C).

### Existing Pathogenicity Predictors Appear to Implicitly Capture the Structural Distance Information

A comparison was done between the two new predictive models: AlphaMissense (AM) + EVE combined, and AM + EVE + Distance. As predictors, AlphaMissense and EVE both showed strongly positive posterior means with credible intervals excluding 0, making them credible predictors of pathogenicity. Distance 1, distance 2, and the distance interaction all have posterior means near 0 with credible intervals that cross 0, indicating no credible effect of distance on predictions. The two models have nearly identical estimates for the shared parameters (Fig 4), which suggests that either distance adds little to the model or that the information coming from AM and EVE dominate the parameter estimates.

**Fig 4.**
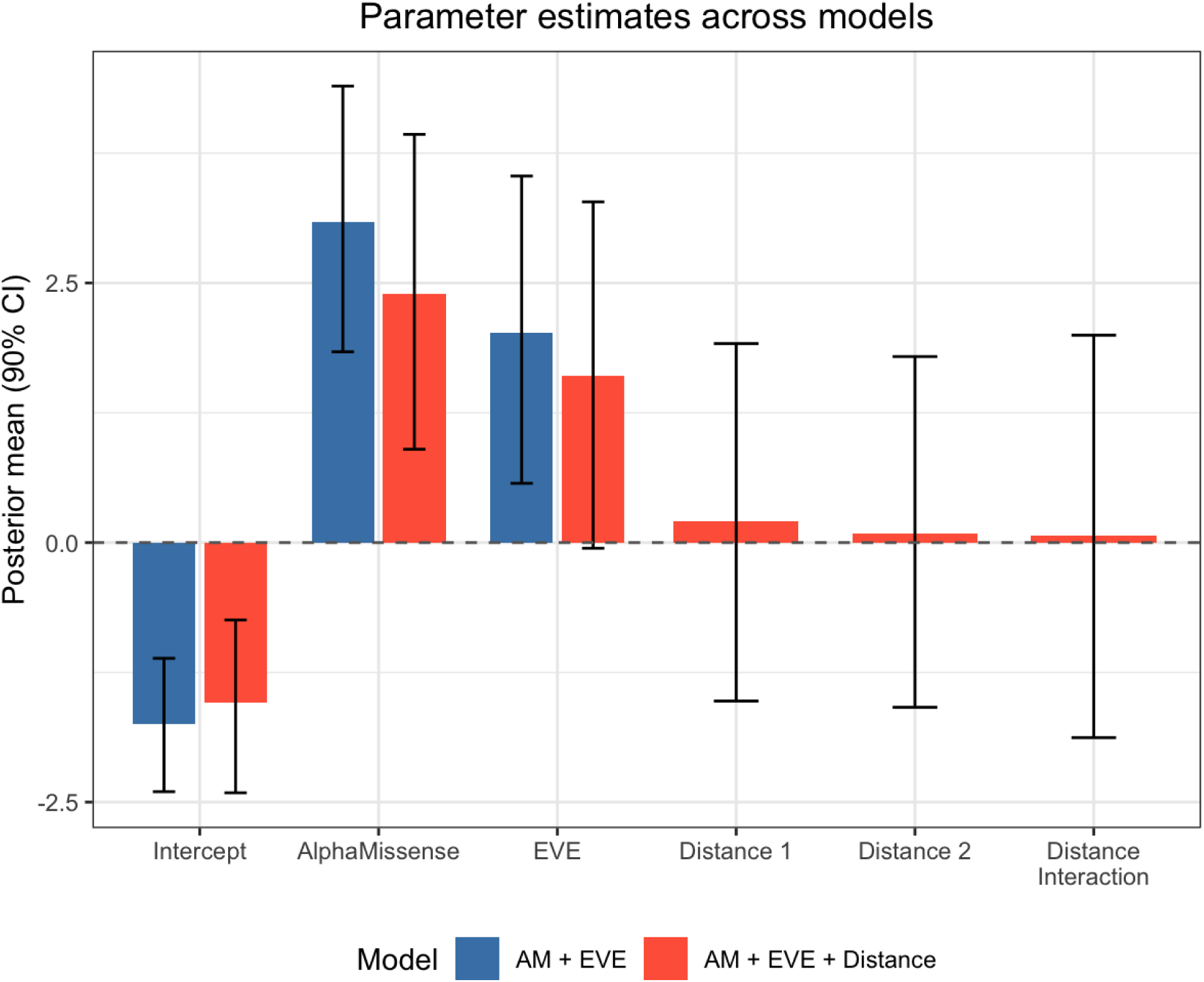
Parameter estimates across models. Posterior mean estimates and 90% credible intervals for the combined AM + EVE model and the AM + EVE + Distance model. Intercept, AlphaMissense, and EVE parameters are shared across both models. Distance 1, Distance 2, and Distance interaction parameters are present only in the extended model. Distance 1 refers to the distance from the variant to the anticodon of the tRNA (nucleotide 35), distance 2 refers to the distance to the 3’ end of the tRNA (last nucleotide), and distance interaction refers to the product of the two distance values.

The existing predictive models, AM and EVE evaluated individually, were compared to the new models: AM and EVE combined, and AM and EVE combined with distance added. When Mean Squared Error (MSE) was evaluated, EVE had the worst performance, combined AM and EVE had the best performance, and when distance was added, MSE was slightly higher than with just AM and EVE alone, indicating a marginally worsened prediction (Table 1).

**Table 1.** Model comparison by mean squared error (MSE) and leave-one-out cross-validation (LOO). ELPD LOO is the expected log pointwise predictive density. p LOO is the effective number of parameters.

| Model | MSE | ELPD LOO | SE | p LOO |
| --- | --- | --- | --- | --- |
| AM | 0.1233 | -42.83 | 4.27 | 1.14 |
| EVE | 0.1427 | -48.40 | 3.88 | 1.19 |
| AM + EVE | 0.1142 | -40.93 | 4.70 | 1.47 |
| AM + EVE + Distance | 0.1152 | -44.88 | 4.05 | 2.19 |

Leave-one-out (LOO) cross-validation was used to evaluate out-of-sample predictive performance, with expected log pointwise predictive density (ELPD LOO) as the primary metric, where less negative values indicate better performance. AM + EVE combined without distance displayed a better performance than when distance was added. However, standard error of ELPD LOO (SE) reveals that the margin of error between the AM + EVE and AM + EVE + Distance models was similar to the difference between the ELPD LOO scores, suggesting the two models are comparable in their ability to generalize to new data. Model complexity is also shown with p LOO, which reveals the effective number of parameters. When distance is added, p LOO jumps to 2.19 compared to 1.47 for just AM + EVE without distance (Table 1).

When comparing the models using AUC calculations for ROC curves, all models performed well, and AM + EVE + distance had the highest AUC, at 0.941. AM + EVE without distance had a slightly lower AUC than AM alone (0.921 vs. 0.923, respectively) (Fig 5).

**Fig 5.**
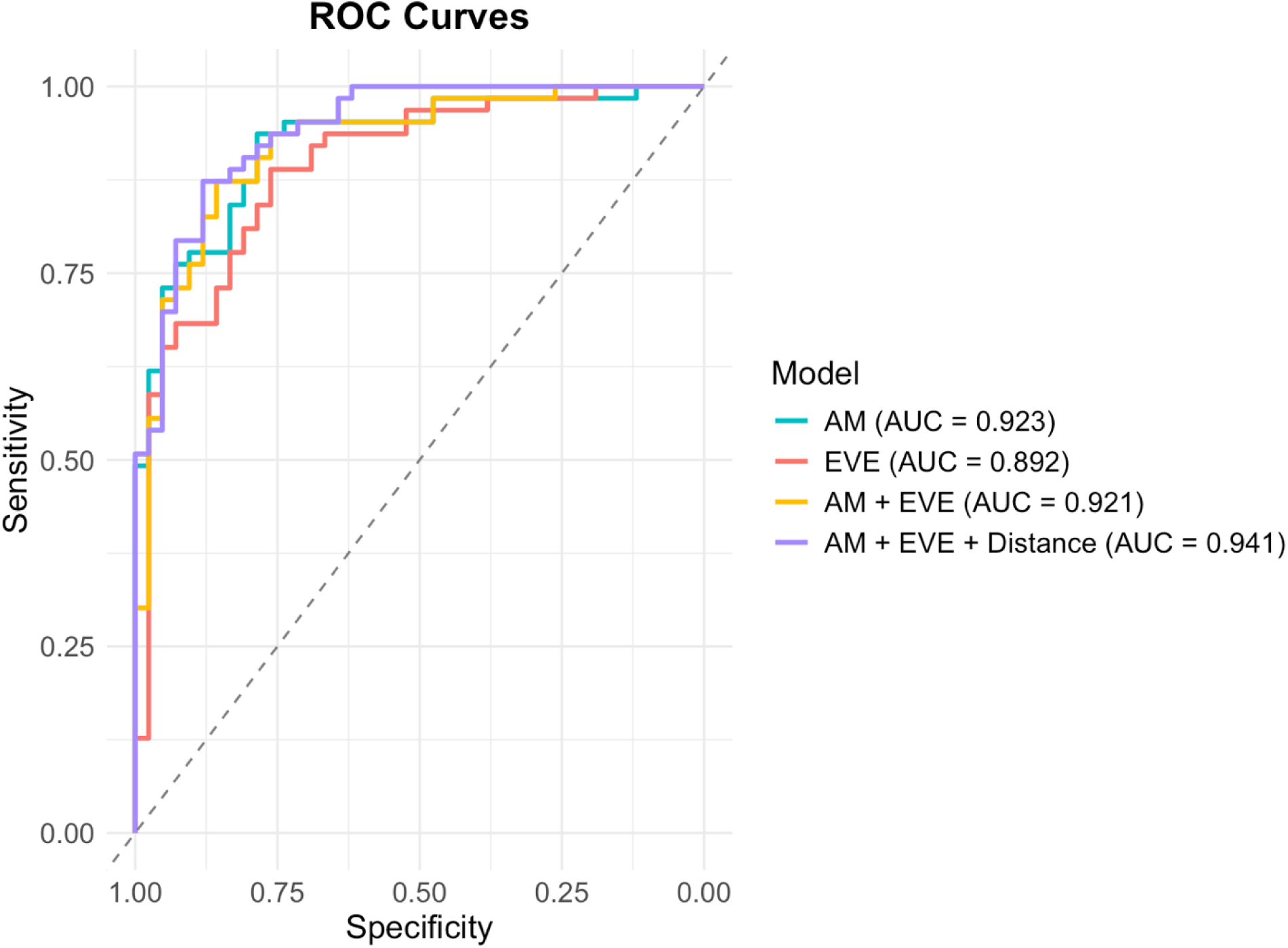
Receiver Operating Characteristic (ROC) curves with Area Under the Curve (AUC) calculations for all four models: AlphaMissense (AM), EVE, AM + EVE, AM + EVE + Distance. Variants were considered benign at scores below 0.30 and pathogenic at scores above 0.70.

Data from Fig 4, Table 1, and Fig 5 all suggests that even though AlphaMissense and EVE do not explicitly encode distance or interface calculations, they do implicitly account for it.

### Investigation of Ambiguous Interface Variants

Pathogenic and benign variants on the interface that are classified as ambiguous by either AlphaMissense or EVE were identified. Ambiguous variants are defined by a pathogenicity score between 0.30 and 0.70. There are more known pathogenic than benign variants in this group, so existing models still miss certain pathogenic variants near the interface (Figs 6 and 7).

**Fig 6.**
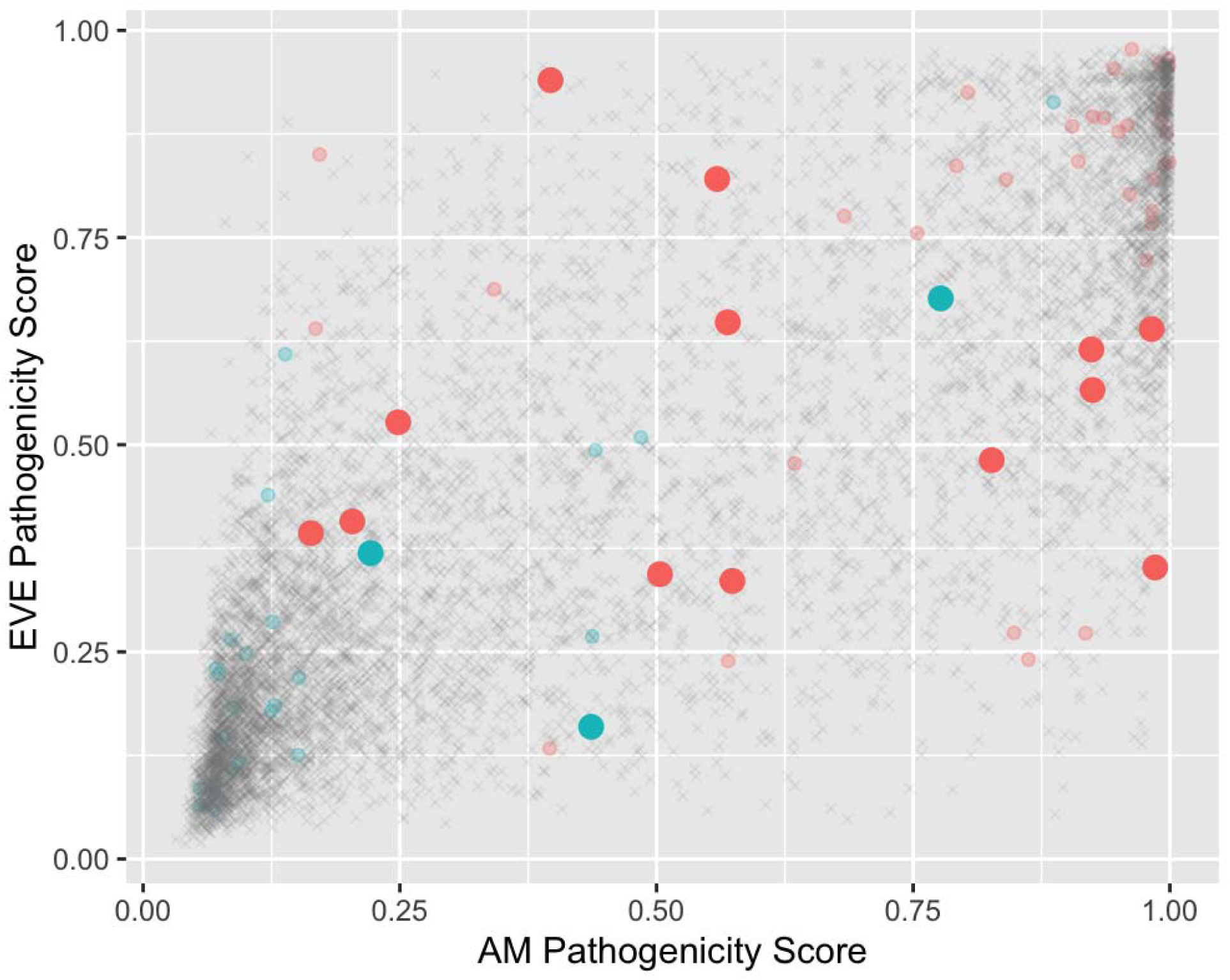
EVE vs. AlphaMissense (AM) pathogenicity scores. Known pathogenic (red) and benign (blue) variants in the interface region are highlighted by using a larger dot.

**Fig 7.**
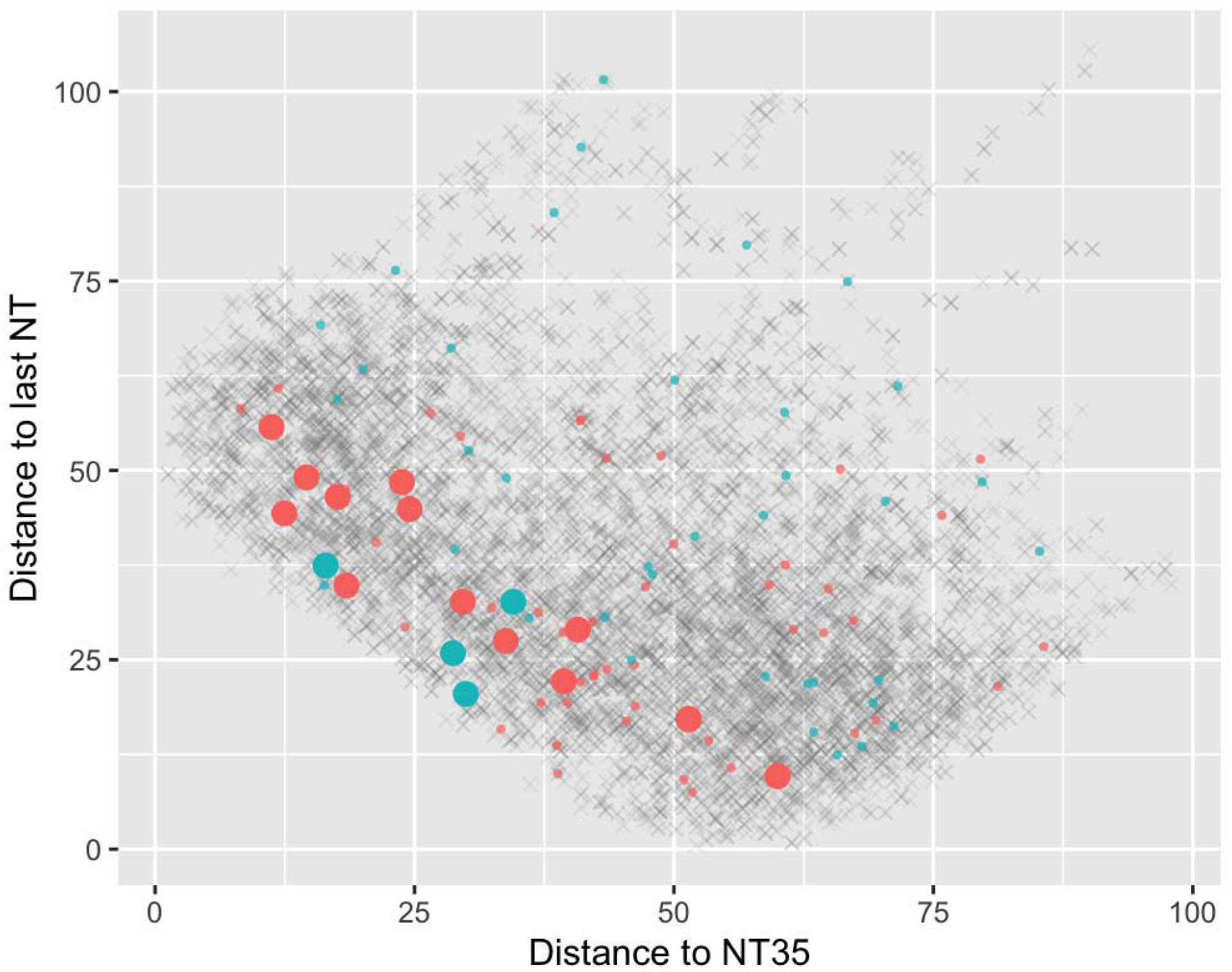
Comparison of mutation distances to the two key nucleotides in angstroms. Known pathogenic (red) and benign (blue) mutations on the interface that are classified with ambiguous pathogenicity (0.30-0.70) by current models (AM + EVE) are highlighted by using a larger dot.

When considering the combined model that includes AlphaMissense and EVE but excludes distance, it is clear that the uncertainty (defined as the width of the Bayesian credible interval) is much higher for variants in the middle range of pathogenicity from roughly 0.30 to 0.70 (Fig 8).

**Fig 8.**
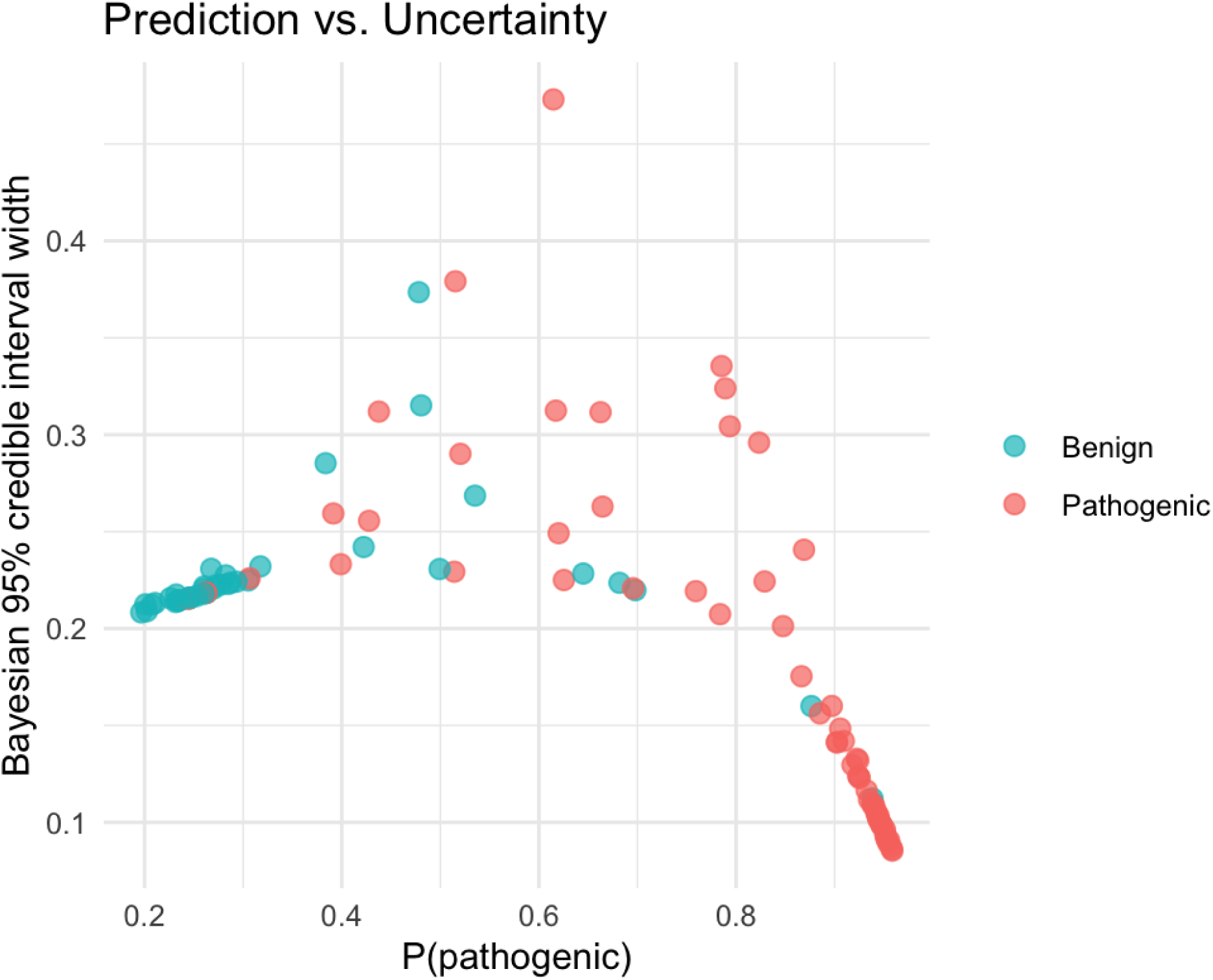
Uncertainty vs. pathogenicity prediction of combined (AM + EVE) model. Uncertainty is measured as the width of the Bayesian 95% credible interval. “True” pathogenicity from UniProt is highlighted in red (pathogenic) and blue (benign).

Given that there relatively few ambiguous variants (17) and thinking that perhaps the AlphaMissense and EVE models were already indirectly incorporating information about distance from the tRNA interface (via higher conservation of those residues), we focused on these 17 variants in order to investigate whether distance alone could provide a way to discriminate among them. We simulated data by sampling with replacement from the original set of these 17 ambiguous variants and generated datasets with 25, 50, 75 and 100 variants. The current dataset with 17 ambiguous variants has a mean AUC around 0.69 with a wide confidence interval. As simulated datasets similar to the ambiguous variants increase in size and the number of ambiguous interface variants approaches 100, the confidence interval narrows and the AUC rises beyond the threshold of good discrimination (Fig 9).

**Fig 9.**
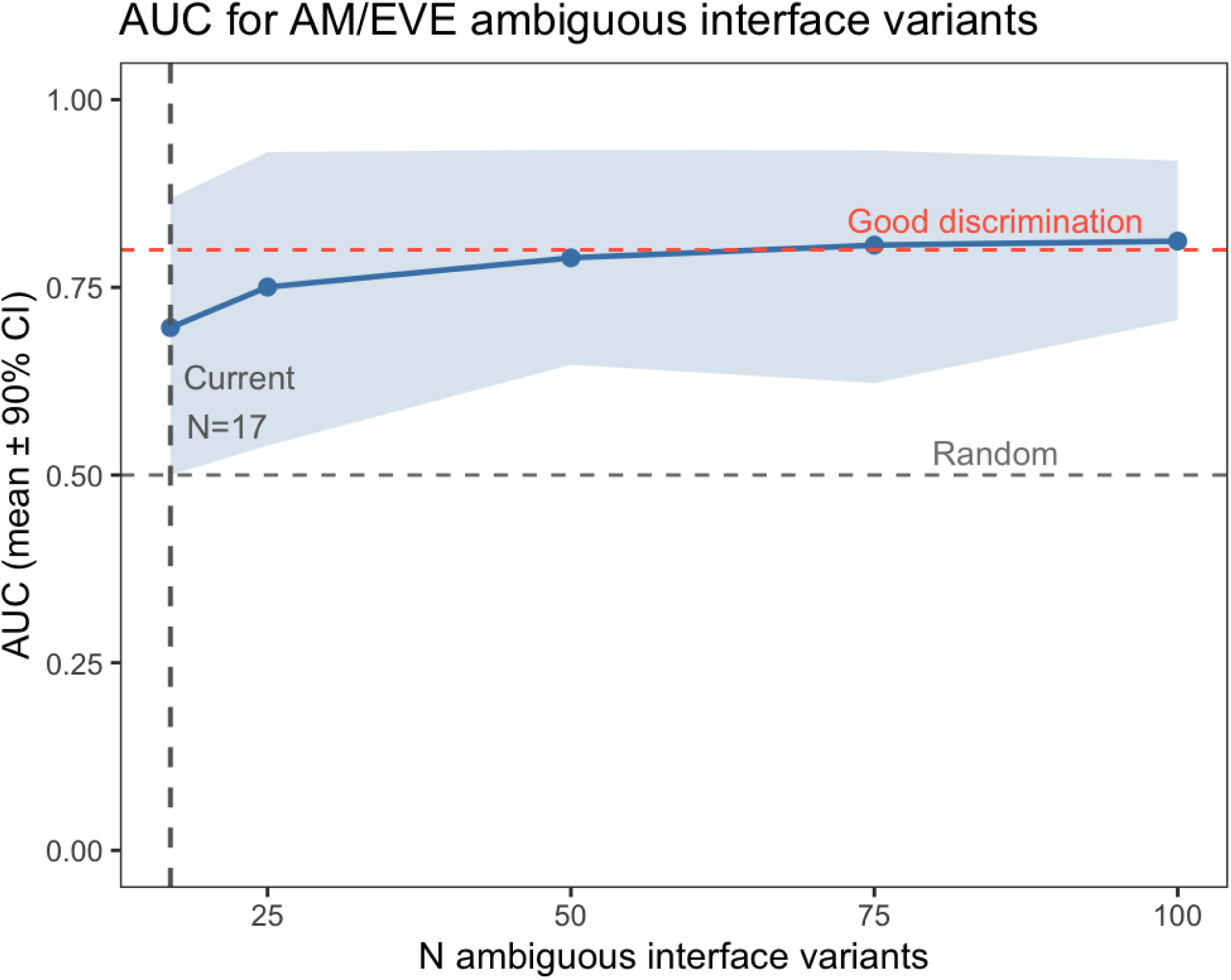
Mean Area Under the Curve (AUC) with 90% confidence intervals for AlphaMissense (AM) and EVE ambiguous interface variants using only distance as a predictor. The dashed lines indicate reference points for non-discriminative (0.5) and good performance (0.8).

## Discussion

### Pathogenic Variants Cluster at the ARS-tRNA Interface

AlphaMissense structures with pathogenicity classifications highlighted revealed that pathogenic variants appear to cluster towards the core of the synthetases near the tRNA-binding region, and benign variants appear to cluster farther out from the core of the synthetases (Fig 1). This was the key observation driving our hypothesis. The tRNA-binding interface is critical for synthetase function, as it is essential for effective recognition of tRNA and aminoacylation (5). Mutations in this region may directly impair the protein’s main function, which would make pathogenicity more likely. Such residues might be expected to show a higher degree of conservation than the hydrophobic core, thereby influencing their weighting in large language models.

The spatial distribution of pathogenic variants in ARSs was found to cluster significantly closer to the tRNA binding interface than that of benign variants (p = 0.0003) (Fig 2B), which suggests that disruption of interactions at this site plays a critical role in determining pathogenic outcomes. Mutations near the interface would directly disrupt the charging of tRNA with their corresponding amino acids, which is essential for protein synthesis. The finding that pathogenic variants cluster near the interface aligns with the idea that mutations near the binding interface could cause disruption of aminoacylation, which could be responsible for some associated disease states.

However, it is important to acknowledge that not all pathogenic ARS variants are expected to cluster near the tRNA binding interface. Dominant mutations associated with CMT disease in homodimeric ARSs may cause disease through gain-of-function or dominant-negative mechanisms involving aberrant heterodimer formation or mutation-induced conformational changes at the dimer interface, rather than through direct disruption of tRNA binding (9). Variants operating through these mechanisms would not necessarily be expected to cluster near the tRNA binding interface, which may account for some of the exceptions observed outside the spatial threshold (Fig 2A). Our ARS-tRNA interface-based approach is therefore likely to be most informative for loss-of-function variants that directly impair aminoacylation and may be less sensitive to gain-of-function or dominant-negative variants whose pathogenicity is mediated by structural mechanisms unrelated to tRNA binding.

This study formally models a spatial threshold defining the binding interface between the synthetases and their corresponding tRNAs (Fig 2A). However, further investigation with more data is needed to define the exact area of the interface more precisely.

An important caveat for this study is that the quality of sequence alignments used have a direct impact on the reliability of the distance calculations. The majority of the structural alignments to create the human ARS-tRNA complexes for distance calculations had fairly high quality, with a range of 75.2–91.7% identity for all synthetases excluding HARS. The sequence alignment for HARS showed notably poor quality with 30.1% matching amino acids (S3 Table). This discrepancy may reflect a use of mismatched isoforms between the structural data used for distance calculations and the reference sequences used by UniProt, EVE, and AlphaMissense. HARS has multiple isoforms, and if the structural model corresponded to a different isoform than the canonical UniProt reference sequence, poor alignment would be expected. Therefore, distance data for HARS should be interpreted with caution. To see if the poor sequence alignment for HARS had a significant effect on the accuracy of the model, we performed an analysis excluding HARS. The analysis resulted in the same qualitative conclusions across all comparisons: significant difference in distribution of pathogenic variants, similar model parameter estimates, and identical conclusions regarding model comparisons.

Furthermore, our overviews of data about pathogenicity of mutations in ARSs revealed that there is a data imbalance between predictive and clinical sources. There is much more predictive information, but these data are less reliable than clinical information. Of the clinical data from UniProt, the majority of the data are classified as neither pathogenic nor benign (S2 Table). This is in part due to a lack of clinical reports of ARS mutations, but it is also influenced by the strict constraints used in this study to classify a mutation as a “true” pathogenic or benign variant.

Additionally, evaluation of existing predictions from EVE and AlphaMissense revealed discrepancies between the two models; AlphaMissense consistently had higher pathogenicity predictions than EVE (S1 Table). The difference in the pathogenicity predictions from these two models likely reflects their unique underlying assumptions about what causes a mutation to be pathogenic that informs their individual approaches.

### New Model Reveals that Existing Models Implicitly Capture Structural Distance Information

#### Data and Parameters Used for the New Model

In order to incorporate the observation of pathogenic variants clustering near the interface into pathogenicity predictions, a Bayesian mixture model was created. The mixture allows us to only apply distance information for the variants that are close enough to the binding site for it to be of significance. A Bayesian model was chosen because it provides uncertainty quantification, allowing for assessment not only of the magnitude of each predictor’s effect but also for an evaluation of the level of confidence in that estimate.

In order to incorporate calculated distances from sites of substitution in the ARSs to the ARS-tRNA interface, raw distance measurements were transformed to a probability scale using the cumulative distribution function (CDF) of a fitted gamma distribution. The gamma distribution of distances to the two key nucleotides shows clustering of most mutations within roughly 100 angstroms (Figs 3A and B, left), which reflects the finite size of the ARS-tRNA interface. The CDF visually supports the hypothesis that pathogenic variants cluster closer to the two key nucleotides than benign variants (Figs 3A and B, right).

Disagreement between AlphaMissense and EVE is most concentrated for variants in the middle range of pathogenicity scores between 0.30 and 0.70 (Fig 3C). The concentration of these ambiguous variants in this intermediate range highlights a fundamental limitation of existing pathogenicity predictions as they do not provide a consistent evaluation of these variants. AlphaMissense has been shown to misclassify a meaningful proportion of clinically validated pathogenic variants, with misclassification rates particularly elevated in intrinsically disordered regions where AlphaFold 2 confidence is low (30). This motivates exploration of whether including explicit structural binding context could resolve these cases where AlphaMissense and EVE are the least reliable.

#### Evaluating the New Model

When comparing all four models, the combined AM + EVE model without distance outperforms either predictor alone in mean standard error (MSE), indicating that the joint predictive power of the two models exceeds that of either model individually (Table 1). However, adding explicit structural distance information to the combined model does not improve performance by MSE or leave-one-out (LOO) cross-validation, as the distance model has a slightly higher MSE than AM + EVE alone, and the difference in expected log pointwise predictive density (ELPD LOO) falls within the standard error (Table 1), indicating the added complexity of the distance model does not translate into significantly better predictions on new data. This is reflected in the effective number of parameters, which increases from 1.47 in the combined AM + EVE model to 2.19 when distance is added (Table 1), confirming that the model gains complexity without a corresponding gain in predictive power by these metrics.

However, when evaluating AUC, the distance model achieves the highest value among all four models (Fig 5). Since the distance model is nested in the AM + EVE model, this difference should be interpreted with caution. LOO cross-validation remains the more informative criterion for evaluating overall model performance.

With the inclusion of interface distance information, posterior mean estimates of AlphaMissense and EVE were both positive, and all forms of distance information evaluated had posterior means near 0 with wide credible intervals (Fig 4), indicating that explicit distance information contributes little beyond what AlphaMissense and EVE already provide. The minimal benefit of explicit distance features supports the idea that existing models already encode structural information, albeit indirectly. Protein language models trained on large sequence databases have been shown to implicitly capture evolutionary, structural, and functional organization across protein space from sequence data alone, without requiring explicit structural input (31). It is also important to note that AlphaFold structures used for our distance calculations are primarily based on sequence evolution data, meaning that structural features extracted from AlphaFold may not provide information truly independent from what AlphaMissense and EVE already capture through multiple sequence alignments (32). AlphaMissense is trained from AlphaFold 2 structures, and EVE uses evolutionary constraints that reflect structural conservation, so it is plausible that these models already implicitly encode the structural information that is added explicitly here with distance to the interface. Interface residues are subject to purifying selection, as mutations at these positions tend to disrupt aminoacylation and are therefore selected against. This selective pressure is reflected in multiple sequence alignments because there is increased conservation of interface positions, meaning that models trained on these alignments may inherently capture proximity to the interface as a consequence of the data on which they were trained.

Overall, we found that AlphaMissense and EVE accurately predict pathogenicity for the majority of variants in the dataset, and because the logistic regression coefficients act globally, they are optimized primarily to fit this well-classified majority. Structural binding distance is most informative for the specific subset of variants where both AlphaMissense and EVE produce ambiguous scores and the variant lies at the tRNA-binding interface, and only 17 of the 105 variants of known pathogenicity (either pathogenic or benign) in the current dataset meet both criteria. With so few ambiguous interface variants, the global likelihood is dominated by the 88 variants for which AlphaMissense and EVE are already informative, which drives the distance parameters toward zero despite a genuine local signal.

### Ambiguous Interface Variants Reveal an Area for Improvement

While explicit structural binding information provided little improvement to existing pathogenicity predictions, AlphaMissense and EVE are shown to have ambiguous pathogenicity scores for several known pathogenic variants located in the ARS-tRNA binding interface (Figs 6 and 7). Additionally, evaluation of uncertainty of the predictions made by the combined AlphaMissense and EVE model excluding explicit distance information revealed that uncertainty is much higher for ambiguous variants with scores between 0.30 and 0.70 (Fig 8). This highlights a gap in current predictive models. More precise characterization of these variants could meaningfully improve the clinical interpretation of interface mutations.

In the current analysis, the sample size of variants on the interface with known pathogenicity but misclassifications from AM and EVE is small. With only 17 ambiguous interface variants, there is a large confidence interval for the AUC when distance is used alone as a predictor (Fig 9), making it difficult to evaluate the utility of incorporating structural binding features in this subset. However, as sample size increases toward 100, the predicted AUC rises beyond the threshold of good discrimination and the confidence interval narrows substantially (Fig 9), suggesting that our approach may become more informative with more data, including additional training set variants and improved structural alignments. This indicates that structural distance is not redundant with existing predictors but is locally informative for a variant class that is currently underrepresented in the dataset. The limiting factor here is the availability of variants of known significance which are ambiguous for both AlphaMissense and EVE. Pathogenic and benign interface variants with confirmed classifications from clinical data are rare in ARS-related diseases. As rare disease sequencing databases expand and variant classification efforts grow, the number of validated interface variants is likely to increase, providing a more powerful test of whether structural binding distance information can improve pathogenicity predictions in this subset of variants.

Beyond sample size limitations, the Euclidean distance calculations used in our model may not be specific enough to correctly identify pathogenicity of variants for which AlphaMissense and EVE have ambiguous classifications. The subset of pathogenic interface variants that are classified as ambiguous by AM and EVE reveals a weak area that existing models using evolutionary and sequence-based information could be improved upon with more granular structural features. While the distance information used here did not provide a significant improvement to existing models, refining the way that structural binding knowledge is incorporated or adding different discriminative features like protein stability could help to improve pathogenicity prediction. In addition, the mixture model acted locally on the interface, but perhaps a more fine-grained model which used distance when AlphaMissense and EVE are ambiguous could be helpful.

Overall, these findings suggest that structurally informed approaches targeting the binding interface remain a promising direction for improving classification of the ambiguous variants that are most difficult to characterize through existing approaches.

### Limitations and Future Directions

It is known that synthetases have non-canonical functions that may involve binding sites other than the ARS-tRNA interface (3), and ARSs also bind with each other to form the multi-synthetase complex (6), but these binding sites were ignored in the model proposed by this study. Recognition of other binding sites could help to improve the accuracy of pathogenicity predictions.

Structural alignment inaccuracies represent a potential source of error. This is particularly relevant given the use of prokaryotic structures, as differences between prokaryotes and eukaryotes could impact alignment quality and make distance calculations less reliable. AlphaFold predictions do not account for ligands, covalent modifications, or environmental conditions, and model protein interactions in a limited way (23), and because our distance calculations are based on ARS structures predicted without their tRNA binding partners, the accuracy of these structural models at the precise interface residues used as distance references may be limited. Furthermore, using two different methods of sequence alignment introduces inconsistency that could further affect the reliability of distance calculations. More high-quality eukaryotic structures of ARS-tRNA complexes would reduce these sources of error. The reliance on static structures also fails to capture how mutations influence protein stability and structural dynamics; molecular dynamics simulations could be explored as a complementary tool to capture conformational changes that a single structural snapshot cannot reflect.

The pathogenicity standards used for validation are also a limitation to be considered. UniProt classifications were used as the primary standard in this study, and they draw heavily from ClinVar submissions, which are known to reflect ascertainment biases toward variants identified in well-studied populations and genes with established disease associations (12). Variants in understudied ARS genes or those identified in underrepresented populations may be systematically underrepresented in these databases, meaning that the true distribution of pathogenic interface variants may differ from what the current dataset reflects. The small sample size of variants with clinically established pathogenicity adds to this limitation, and this could be addressed by incorporating additional cytoplasmic ARS data as it emerges, extending the model to synthetases currently excluded due to data limitations, expanding to mitochondrial synthetases, or even applying this framework to other nucleic acid binding proteins.

Finally, two directions emerge from this work. First, adding another structural feature in addition to distance could support discrimination between pathogenic and benign variants. Second, because the existing models seem to be able to perform well on most of the variants, rather than applying distance as a predictor on the interface regime, a model that applies distance only when AlphaMissense and EVE scores are ambiguous could be more informative. These two directions together point toward a hybrid framework in which large-scale AI models such AlphaMissense and EVE handle the majority of variants confidently, while a more structurally informed local model fills the gaps for the subset of variants which are ambiguous for these tools.

## Methods

### Data compilation

Mutations for the cytoplasmic ARSs were compiled from UniProt. Due to the large volume of data, bulk protein data were first downloaded to the Vermont Advanced Computing Center (VACC), from which specific synthetase data were then extracted and downloaded locally for analysis.

Pathogenicity annotations in UniProt were used as the standard for “true” pathogenicity in this model. For the UniProt data, pathogenic variants were labeled with a pathogenicity of 1, and benign variants were labeled with a pathogenicity of 0. UniProt annotations were classified as pathogenic if the UniProt annotation contains “Pathogenic” and/or “Likely Pathogenic” and classified as benign if the UniProt annotation contains “Benign” and/or “Likely Benign.” This method ignored individual mutations with conflicting or uncertain pathogenicity annotations in UniProt. Because UniProt entries include combinations of different data sources, there is often conflicting information for a single variant. Mutations with any conflicting information in UniProt were ignored so that the training data set from UniProt with “true” pathogenic and benign variants strictly included mutations with strong clinical evidence of pathogenicity. However, this caused our training data set to exclude certain known pathogenic variants. For example, the Y454S substitution mutation in HARS has been shown to be linked to Type IIB Usher syndrome, which often presents with symptoms including childhood deafness, blindness, and episodic hallucinations during acute illness (33). Our classification in the data downloaded from UniProt is “Pathogenic, Variant of uncertain significance,” so it does not get incorporated into the data we use because it has conflicting interpretations of pathogenicity.

AlphaMissense and EVE pathogenicity scores were also downloaded using the VACC and subsequently extracted for the synthetases of interest. The EVE, AlphaMissense, and UniProt data were then merged into a larger data set containing any mutations with pathogenicity information from at least one of the three sources.

### Alignments

ARS structures were aligned to their corresponding tRNA molecules using protein structures generated from AlphaFold. Two different methods were used depending on availability of structures. For DARS, GARS, IARS, QARS, RARS, SARS, TARS, VARS, WARS, YARS, and HARS, a prokaryotic version of the bound structure with the ARS in complex with its corresponding tRNA was available, so the human synthetase structure was aligned to the prokaryotic synthetase in PyMOL (34). Structures with extremely poor structural alignments to prokaryotic synthetases (IARS and SARS) were excluded from analysis. For CARS, KARS, LARS, and NARS, the AlphaFold server (22) was used to align the human FASTA sequence for the synthetase to the human tRNA sequence from RNAcentral (35).

We wrote a program in R (36) that calculated minimum Euclidean distances from each residue on the protein to each nucleotide on the tRNA molecule in angstroms. It should be noted that due to differences in crystal structures, nucleotide sequence lengths of tRNAs varied from 72-83 in our dataset. While AlphaFold predictions are highly accurate in many cases, they can differ from experimental structures, particularly in side-chain conformations and regions involving ligand interactions not included in the prediction (23). The distance calculations in this study should therefore be interpreted with this limitation in mind.

UniProt, AlphaMissense and EVE use a coordinate system based on the UniProt reference sequence, which in some cases differed from the sequences used in our structural alignments for tRNA binding site distance calculations. This meant position numbers were not directly comparable across data sources. For each enzyme, we used the AlignSeqs function in the DECIPHER package in R (37) to align the reference sequence to its corresponding structural sequence. All data was then merged into a single data frame.

### Bayesian model

Full description of the model can be found in the supplementary information. Briefly, a Bayesian mixture model was developed to integrate existing pathogenicity scores from AlphaMissense and EVE with structural distances to the two key catalytic nucleotides of the tRNA, the anticodon (d_1_) and the 3’ end (d_2_). The model partitions variants into two structural regimes, interface and non-interface, defined by a probabilistic threshold U on the sum of distances to the two key nucleotides (d_1+_ d_2_): the interface regime region (d_1+_ d_2_ ≤ U) contains mutations close to the key nucleotides, and the non-interface region (d_1+_ d_2_ > U) contains mutations that are farther away. Rather than applying a hard cutoff, U is treated as a latent variable averaged over equally weighted K discrete values, hence allowing the threshold itself to also be a parameter to estimate. The threshold is represented as a diagonal line in the two-dimensional space defined by the distances to the anticodon (nucleotide 35) and the 3’ terminal nucleotide of the tRNA (see for example the diagonal line in Fig 2).

For variants assigned to the interface regime (distanceSum = *d*_1_ + *d*_2_ ≤ U), the likelihood model uses all five covariates to predict pathogenicity,

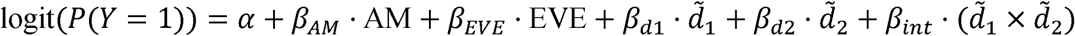

Here, AM and EVE are the AlphaMissense and EVE pathogenicity scores respectively, *d̃*_1_and *d̃*_2_ are gamma cumulative distribution function (CDF) transformations of *d̃*_1_and *d̃*_2_ respectively. *d̃*_1_ *x d̃*_2_ captures the joint proximity interaction. The CDF transformation expresses each distance relative to the reference distribution of all residues, such that low values indicate unusually close proximity. The interface regime parameters (*α, β_AM_, β_EVE_, β_d1_, β_d2_*, and *β_int_*) are informed by AM, EVE, and gamma CDF-transformed structural distance measurements.

For variants assigned to the non-interface regime (distanceSum > U), distance information is omitted because pathogenicity in this regime is not expected to be mediated by proximity to the tRNA binding interface. Pathogenicity is instead modeled using AM and EVE scores alone:

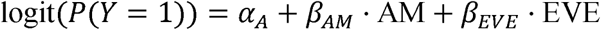

The non-interface regime parameters *(α_A_, β_AM_, β_EVE_*) are informed by an AM + EVE only model fitted to the full dataset. All model parameters are estimated using Hamiltonian Monte Carlo as implemented in the probabilistic programming language Stan (38), with weakly informative priors. Missing EVE scores are handled separately through probabilistic imputation, in which missing values are estimated from the observed AlphaMissense score, using the correlation between the two predictors.

### Model performance evaluation

We evaluated the models using the following three methods:

- MSE: measures the averaged square difference between the estimated probabilities and the observed outcomes.
- LOO: leave-one-out cross validation evaluates how well the model will perform on new data by running the model on the entire dataset minus one observation, doing this once with each observation absent, and estimating the expected log predictive density (ELPD) evaluated as: 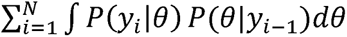 where *P(θ*|*y_i_*_-1_) is the posterior distribution of *θ* given the dataset without the *i*’th observation, and *P(y*_i_|θ) is the probability of the *i*’th observation given the parameters.
- AUC (area under the curve): probability that a pathogenic variant will be ranked higher than a benign one. This provides a way to measure how well the model discriminates between pathogenic and benign.

All programming was done in the R programming language (version 4.4.1) (36), with code available in Zenodo (DOI: 10.5281/zenodo.20124785).

Claude (39) was used to assist with statistical analysis code development, Stan model implementation, R scripting, and for providing manuscript feedback. All analyses were completed, interpreted, and verified by the authors. All scientific conclusions are the sole responsibility of the authors.

## Supporting information

Bayesian Model Development and Assessment

## Acknowledgements

We deeply thank Michael Betancourt for his advice and support on Bayesian statistical modeling and workflow. We thank Jean Gabriel Young and Peter Callas for helpful discussions on the model approach, and special thanks to Becky Tang for going through the supplement describing the Bayesian model.

We also thank Charles Carter for valuable discussions regarding statistical approaches, and Julie Dragon for helpful input and feedback during the development of this project.

We would also like to thank Bryan Ballif and Alicia Ebert for their feedback and advice on this work.

The authors acknowledge the Vermont Advanced Computing Center (VACC) at the University of Vermont for providing computational resources that have contributed to the research results reported within this paper.

## Supporting Information Captions

**S1 Table.** Comparison of data sources. “Unique Proteins” shows how many synthetases are included. “Total Mutations” for UniProt includes variants downloaded from UniProt that were not included in the final analysis due to either a lack of pathogenicity annotation or conflicting annotations; see S2 Table for more detail. “Average Pathogenicity Score” is shown for EVE and AlphaMissense (AM), but not for UniProt or Distance because they do not have quantitative values assigned for pathogenicity of mutations.

**S2 Table.** Mutation data summary. Individual proteins and corresponding variant counts for which structural distance information was available are included. The number of mutations with an AlphaMissense (AM) and EVE score are listed, but EVE lacks data for NARS. UniProt annotations (pathogenic and benign) are also included. “With UniProt annotation” includes any variant with an annotation on UniProt under “clinical significance.” Some annotations contained conflicting interpretations of pathogenicity, so “Pathogenic (UniProt)” and “Benign (UniProt)” show the variants with consistent annotations on UniProt; these were the only variants included in the “true” pathogenicity classifications. See methods for how annotations were classified as pathogenic or benign.

**S3 Table.** Summary of Sequence Alignment Quality. Comparing aligned distance data sequences to UniProt data sequences. “Total Rows” indicates the number of variants available in the dataset for each protein (this includes only variants that exist both in UniProt/AlphaMissense and in the pdb sequence used for distance calculations); these can include multiple variants in the same position with different substitutions. “Available AA Pairs” represents the number of amino acids paired for each sequence alignment (this includes only one amino acid per position). “Matching AAs” shows the number of amino acid pairs in each sequence alignment that were identical. “Percent Matching” shows the ratio of matching amino acids to total amino acid pairs available.

