## Supplementary material for "Structural distance at the tRNA synthetase active site interface predicts pathogenicity but is captured by AlphaMissense and EVE except among score-ambiguous variants": Bayesian Model Development and Assessment

Bayesian Prediction of Pathogenic Variants in tRNA Synthetases using Existing Models and Structural Information

### Supplementary Information: Bayesian Model Development and Assessment

Ramiro Barrantes Reynolds*

May 15, 2026

#### Contents


**1 Model** **S2**

1.1 Overview . . . . . S2

1.2 Data and Notation . . . . . S2

1.3 Distance Transformation . . . . . S3

1.4 Assignment to Interface vs Non-Interface Regimes . . . . . S5

1.5 Linear Predictors . . . . . S5

1.6 Likelihood . . . . . S6

1.7 Priors . . . . . S6

1.8 Posterior Probability of Pathogenicity . . . . . S7

1.9 Model Parameters Summary . . . . . S7

1.10 Notes on Prior Choices and Model Implementation . . . . . S7

**2 Prior Predictive Checks** **S8**

**3 Posterior Predictive Checks** **S9**

3.1 Other Model Diagnostics . . . . . S11

**4 Model Assessment** **S12**

4.1 Computational Faithfulness . . . . . S13

4.2 Simulation Based Calibration . . . . . S13

**5 Model Sensitivity** **S14**

**6 Comparison with Other Models** **S16**

**7 Power Analysis** **S18**

**8 Description and Validation of Missing Data Imputation Model** **S18**
8.1 Model . . . . . . . . . . . . . . . . . . . . . . . . . . . . . . . . . . . . . . S19
8.1.1 Prior Predictive Checks . . . . . . . . . . . . . . . . . . . . . . . . . . S20
8.2 Posterior Predictive Checks . . . . . . . . . . . . . . . . . . . . . . . . . . . S22
8.3 Final Imputed Observations . . . . . . . . . . . . . . . . . . . . . . . . . . . S24

**9 Session Information** **S25**

### 1 Model

#### 1.1 Overview

We would like to predict pathogenicity of a position in an enzyme. We have a set of positions for which we know if an enzyme is pathogenic $\{y_i = 1\}$ or not, $\{y_i = 0\}$. We also know, based on the biology of the enzymes, that there are two key residues (which we call key residues 1 and 2), and we hypothesize that pathogenicity is influenced by proximity to one or both of these residues, which also corresponds to an area of the enzyme called the *interface*.

We also have two "black box" measurements predictive of pathogenicity: AlphaMissense, $X_a$ [Tordai et al., 2024], and Eve, $X_b$ [Frazer et al., 2021].

We model the pathogenicity of protein variants using a Bayesian logistic regression based on these four measurements: AlphaMissense, Eve, distance to key residue1 and distance to key residue 2. Based on a threshold the model separates the enzyme into two parts: *interface*, if the variant is spatially close to both functional residues within a region determined by the sum of the distances to both residues; and *non-interface* otherwise. On the *interface* regime the model relies on all four measurements, while on the *non-interface* regime it relies only on the two black box measurements, as we hypothesize that beyond a certain distance, the key residues will not have an impact. The threshold is treated as a discrete latent variable and we marginalize over it to obtain a threshold-averaged predicted probability of pathogenicity for each variant (although we also report the threshold value with the highest posterior probability, which makes sense biologically).

#### 1.2 Data and Notation

Let $n = 1, \ldots, N$ index variants. For each variant we observe:

- $y_n \in \{0, 1\}$: pathogenicity label (1 = pathogenic, 0 = benign), observed for a subset of variants;
- $\mathrm{AM}_n \in \mathbb{R}$: AlphaMissense pathogenicity score;
- $\mathrm{EVE}_n \in \mathbb{R}$: EVE score;
- $d_{1n}, d_{2n} > 0$: distance from the variant to key residues 1 and 2, respectively.

Table S1 shows the list of data point that we have for each position in the enzyme. For this analysis, we considered all positions of unknown pathogenicity that had all four measurements (7212 positions). Since we wanted to consider all positions with pathogenic information (105), we developed a Bayesian model to impute 29 positions that did not have EVE measurements (see Imputation section).

| Name | Value |
|---|---|
| Total Variants | 9391 |
| Number of AM Scores | 9391 |
| Number of EVE Scores | 7212 |
| Number of distance to key residue1 | 9391 |
| Number of distance to key residue2 | 9391 |
| Number of Pathogenicity Labels | 105 |
| Number of Pathogenic Variants | 63 |
| Number of Benign Variants | 42 |
| Number of AM Scores with Pathogenicity Labels | 105 |
| Number of EVE Scores with Pathogenicity Labels | 76 |

**Table S1:** Summary of available data for model covariates and outcomes

We also specify $K$ candidate threshold values $U_1 < U_2 < \cdots < U_K$ with associated prior weights $\boldsymbol{\omega} = (\omega_1, \ldots, \omega_K)$, $\sum_k \omega_k = 1$. For this work, we set $\omega_k = \frac{1}{K}$ for all $k$, although one could consider learning these weights from the data or through biological priors.

##### 1.3 Distance Transformation

Raw distances are mapped to a probability scale using the cumulative distribution function (CDF) of a Gamma distribution with fixed hyperparameters $\alpha_j$ and $\beta_j$:

$$\tilde{d}_{jn} = F_\Gamma(d_{jn} \mid \alpha_j, \beta_j), \quad j \in \{1, 2\}, \tag{S1}$$

where $F_\Gamma(\cdot)$ denotes the Gamma CDF. The Gamma-CDF maps distances to $[0, 1]$: where $\tilde{d}_{jn}$ indicates the probability of observing a distance less than or equal to $d_{jn}$ under the reference Gamma distribution. The closer the distance is to zero, the smaller the CDF, indicating a lower probability of observing such a close distance under the reference distribution (i.e. under the null). Conversely, larger distances will have CDF values closer to one, indicating a higher probability of observing such distances under the reference distribution. This transformation allows us to capture the intuition that variants closer to key residues are more likely to be pathogenic. We will also add an interaction term, to capture the intuition that being on the interface may have a different effect than being close to each one independently.

We centered the transformed distances to improve computational inference.

$$\tilde{d}_{jn}^{\text{centered}} = \tilde{d}_{jn} - \frac{1}{N}\sum_{i=1}^{N}\tilde{d}_{ji} \tag{S2}$$

The fixed hyperparameters $\alpha_j$ and $\beta_j$ come from fitting all the data to a gamma distribution. Figure S1 shows the fitted gamma distributions for the sums of all the distances (left) and for distances corresponding to pathogenic residues (right), as well as a 2D plot of the distances to the two key residues and their distribution.

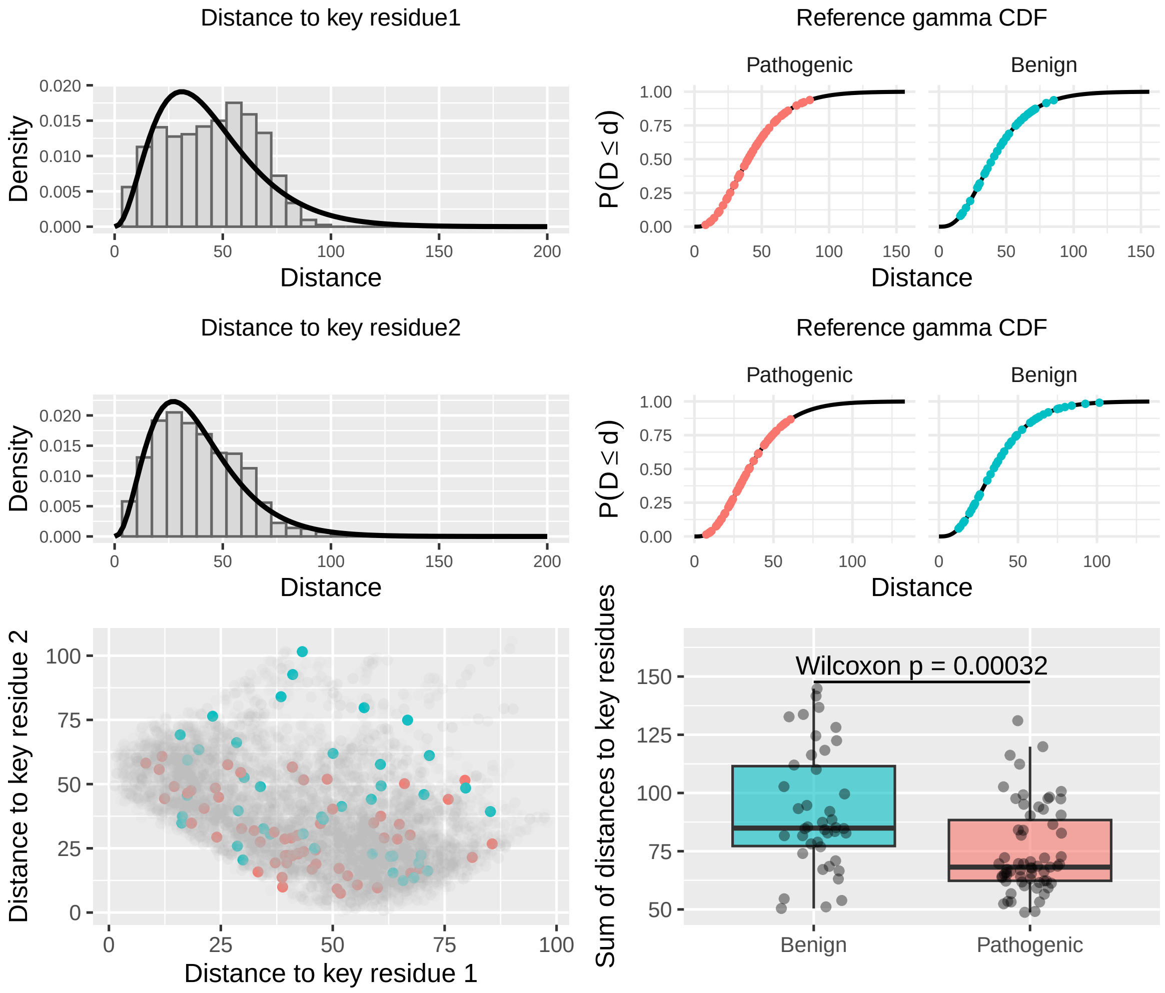


**Figure S1:** Reference gamma distribution for the distance to key residue1 (top row), key residue2 (middle row) and scatter plot comparing both (bottom row). The gamma CDFs show the pathogenic (red) and benign (blue) variants.

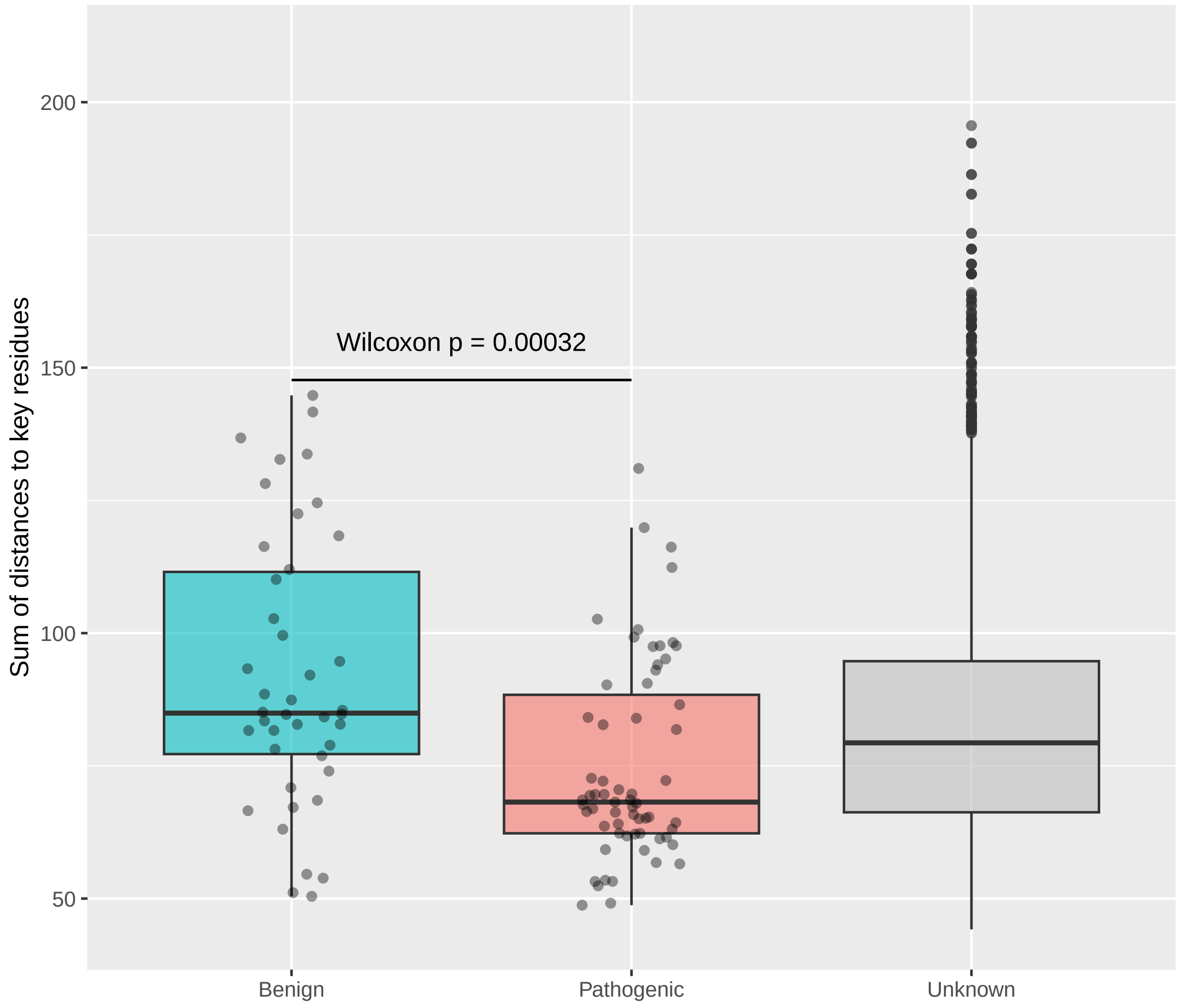


**Figure S2:** Distribution of the sums of distances to the two key residues for the variants known to be Benign, Pathogenic and all the rest (Unknown).

##### 1.4 Assignment to Interface vs Non-Interface Regimes

For a given threshold $U_k$, variant $n$ is assigned to the *interface* if the sum of the distances is below a threshold:

$$z_{nk} = \mathbf{1}[d_{1n} + d_{2n} \leq U_k]\,. \tag{S3}$$

This AND condition expresses the biological hypothesis that joint spatial proximity to both key residues influences pathogenicity if they lie along a region of the protein.

##### 1.5 Linear Predictors

**Interface regime ($z_{nk} = 1$).**

$$\eta_{nk}^{(\text{prox})} = \alpha + \text{AM}_n \cdot \beta_{\text{AM}} + \text{EVE}_n \cdot \beta_{\text{EVE}} + \tilde{d}_{1n} \cdot \beta_{d_1} + \tilde{d}_{2n} \cdot \beta_{d_2} + \tilde{d}_{1n}\,\tilde{d}_{2n} \cdot \beta_{\text{int}}, \tag{S4}$$

where $\beta_{d_1}$ and $\beta_{d_2}$ capture the independent effects of proximity to each key residue, and $\beta_{\text{int}}$ captures their interaction.

**Non-interface regime ($z_{nk} = 0$).**

$$\eta_{nk}^{(\text{away})} = \alpha^A + \text{AM}_n \cdot \beta_{\text{AM}}^A + \text{EVE}_n \cdot \beta_{\text{EVE}}^A. \tag{S5}$$

Variants far from both key residues are scored by a reduced model that relies only on the AM and EVE predictors, with a separate intercept and separate regression coefficients to allow the relationship between scores and pathogenicity to differ across regimes.

#### 1.6 Likelihood

The threshold $U_k$ is unknown and is treated as a discrete latent variable. We marginalize over it using the mixture weights $\omega_k = \frac{1}{K}$. For observed variants the marginal log-likelihood is:

$$\log p(y_n \mid \boldsymbol{\theta}) = \log \sum_{k=1}^{K} \omega_k \, p(y_n \mid \eta_{nk}^{(z_{nk})}) \tag{S6}$$

where

$$p(y_n \mid \eta_n) = \text{Bernoulli-logit}(\eta_n) = \sigma(\eta_n)^{y_n} \left(1 - \sigma(\eta_n)\right)^{1-y_n}, \quad \sigma(\eta_n) = \frac{1}{1 + e^{-\eta_n}} \tag{S7}$$

and $\eta_{nk}^{(z_{nk})}$ denotes the linear predictor for variant $n$ under threshold $k$, selecting the appropriate *interface* or *non-interface* regime.
Variants with missing $y_n$ are inferred via posterior predictive simulation.

#### 1.7 Priors

All regression parameters receive weakly informative normal priors with user-specified standard deviations:

$$\alpha \sim \mathcal{N}(0,\, \sigma_\alpha^2) \tag{S8}$$

$$\beta_{\text{AM}} \sim \mathcal{N}(0,\, \sigma_{\text{AM}}^2) \tag{S9}$$

$$\beta_{\text{EVE}} \sim \mathcal{N}(0,\, \sigma_{\text{EVE}}^2) \tag{S10}$$

$$\beta_{d_1} \sim \mathcal{N}(0,\, \sigma_{d_1}^2) \tag{S11}$$

$$\beta_{d_2} \sim \mathcal{N}(0,\, \sigma_{d_2}^2) \tag{S12}$$

$$\beta_{\text{int}} \sim \mathcal{N}(0,\, \sigma_{\text{int}}^2) \tag{S13}$$

$$\alpha^A \sim \mathcal{N}(0, \sigma_{\alpha^A}^2) \tag{S14}$$

$$\beta_{\text{AM}}^A \sim \mathcal{N}(0,\, \sigma_{\text{AM}}^2) \tag{S15}$$

$$\beta_{\text{EVE}}^A \sim \mathcal{N}(0,\, \sigma_{\text{EVE}}^2). \tag{S16}$$

The following are the values for the standard deviations of the priors:

$$\sigma_{\alpha} = 1.2 \tag{S17}$$

$$\sigma_{\text{AM}} = 1.2 \tag{S18}$$

$$\sigma_{\text{EVE}} = 1.2 \tag{S19}$$

$$\sigma_{d_1} = 1.2 \tag{S20}$$

$$\sigma_{d_2} = 1.2 \tag{S21}$$

$$\sigma_{\text{int}} = 1.2 \tag{S22}$$

$$\sigma_{\alpha^A} = 1.3 \tag{S23}$$

$$\sigma_{\text{AM}_A} = 1.3 \tag{S24}$$

$$\sigma_{\text{EVE}_A} = 1.3 \tag{S25}$$

##### 1.8 Posterior Probability of Pathogenicity

Given observed data $\mathbf{y}_{\text{obs}}$ and model parameters $\boldsymbol{\theta} = \{\alpha, \beta_{\text{AM}}, \beta_{\text{EVE}}, \beta_{d_1}, \beta_{d_2}, \beta_{\text{int}}, \alpha^A, \beta^A_{\text{AM}}, \beta^A_{\text{EVE}}\}$, the posterior distribution over parameters is:

$$p(\boldsymbol{\theta} \mid \mathbf{y}_{\text{obs}}) = \frac{p(\mathbf{y}_{\text{obs}} \mid \boldsymbol{\theta})\, p(\boldsymbol{\theta})}{p(\mathbf{y}_{\text{obs}})} \tag{S26}$$

$$p(\boldsymbol{\theta} \mid \mathbf{y}_{\text{obs}}) \propto p(\mathbf{y}_{\text{obs}} \mid \boldsymbol{\theta})\, p(\boldsymbol{\theta}) \tag{S27}$$

$$p(\boldsymbol{\theta} \mid \mathbf{y}_{\text{obs}}) \propto \underbrace{\prod_{i=1}^{N_{\text{obs}}} \sum_{k=1}^{K} \omega_k\, p(y_i \mid \eta_{ik}^{(z_{ik})})}_{\text{likelihood}} \underbrace{p(\boldsymbol{\theta})}_{\text{prior}} \tag{S28}$$

##### 1.9 Model Parameters Summary

| Parameter | Description | Regime |
|---|---|---|
| $\alpha$ | Intercept | *interface* |
| $\beta_{\text{AM}}$ | AlphaMissense effect | *interface* |
| $\beta_{\text{EVE}}$ | EVE score effect | *interface* |
| $\beta_{d_1}$ | CDF distance to residue 1 effect | *interface* |
| $\beta_{d_2}$ | CDF distance to residue 2 effect | *interface* |
| $\beta_{\text{int}}$ | Distance interaction effect | *interface* |
| $\alpha^A$ | Intercept | *non-interface* |
| $\beta^A_{\text{AM}}$ | AlphaMissense effect | *non-interface* |
| $\beta^A_{\text{EVE}}$ | EVE score effect | *non-interface* |

##### 1.10 Notes on Prior Choices and Model Implementation

We used visual inspection of simulated parameters as well as prior predictive checks to select priors.

We also had 29 missing observations for EVE scores which had observed pathogenicity labels. We used an imputation method to infer those. See section 8.

Finally, we made two key implementation choices when doing inference on the real data:

- Given that we only had 105 observation with pathogenicity information, inference on the non-interface model became more uncertain as $U$ increased. Therefore, we decided to use fixed parameters for the non-interface regime when doing inference on the real data rather than trying to infer it. These fixed parameters were obtained from fitting the reduced model on all the data.
- As mentioned, we average over the discrete threshold variable $U_k$ rather than treating it as a parameter to be inferred.

### 2 Prior Predictive Checks

If we denote $\theta = \{\alpha, \beta_{\text{AM}}, \beta_{\text{EVE}}, \beta_{d_1}, \beta_{d_2}, \beta_{\text{int}}, \alpha^A, \beta^A_{\text{AM}}, \beta_{\text{EVE}}\}$ we can now generate simulated parameters:

$$\tilde{\theta} \sim p(\theta)$$

And we can use those simulated parameters to generate simulated observations :

$$y \sim P(y|\tilde{\theta})$$

We can check that the distribution of $\tilde{\theta}$ and any relevant summary functions reflect our domain expertise.

We removed any datasets which had zero pathogenic or zero benign observations, as those datasets would not be useful for our inference and would introduce degeneracies in the model fitting.

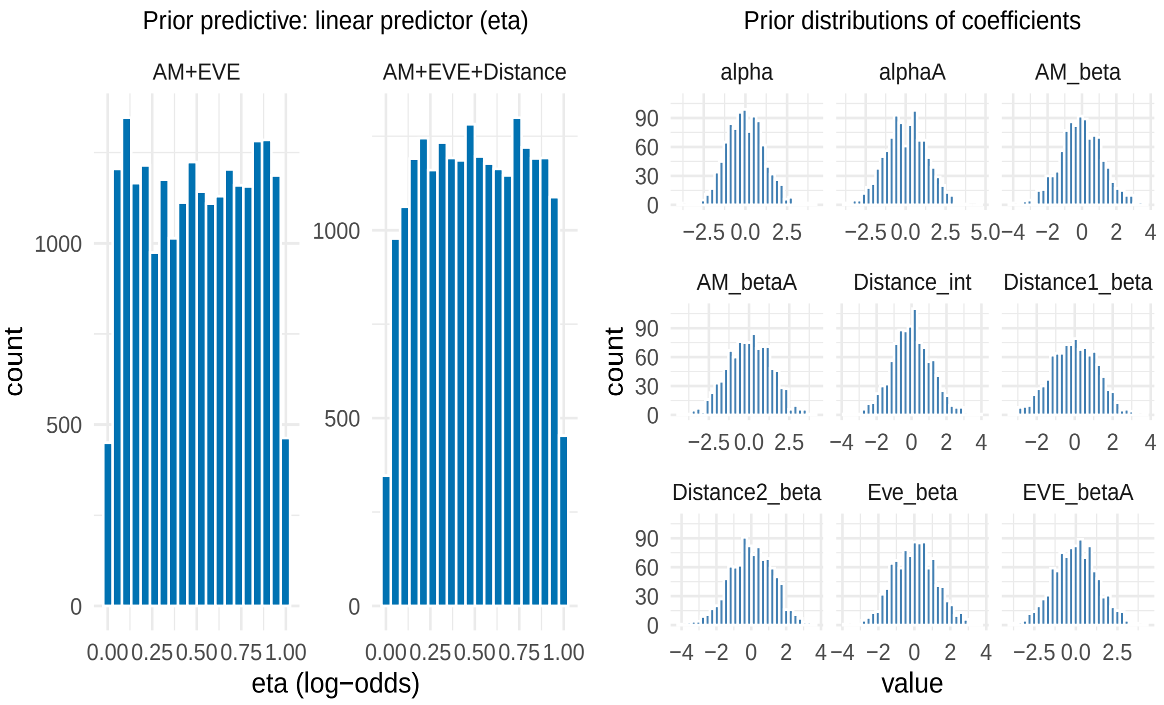


**Figure S3:** Prior distributions from simulated data for model that considers a full model with AM and EVE and distance data. Distribution of predicted probabilities should favor too much either extreme (0 or 1), indicating that the priors are too wide (left graph). The prior distributions for the coefficients are shown on the upper right graph.

### 3 Posterior Predictive Checks

Once we have verified the relevance of the priors, we will check how well does the model reflect real data. This we do using posterior predictive checks, where we fit our real data ($\{y'\}$) and obtain the posterior distribution for the parameters, which we call $\theta'$:

$$\theta' \sim p(\theta|y')$$

and then generate observations from the posterior parameters:

$$\tilde{y} \sim p(y|\theta')$$

We can then compare the generated observations and the real data using relevant summary statistics. We first looked at the average number of pathogenic variants obtained from the generated observations,the actual number of pathogenic variants lies within what is reasonable (Figures **??** and **??**).

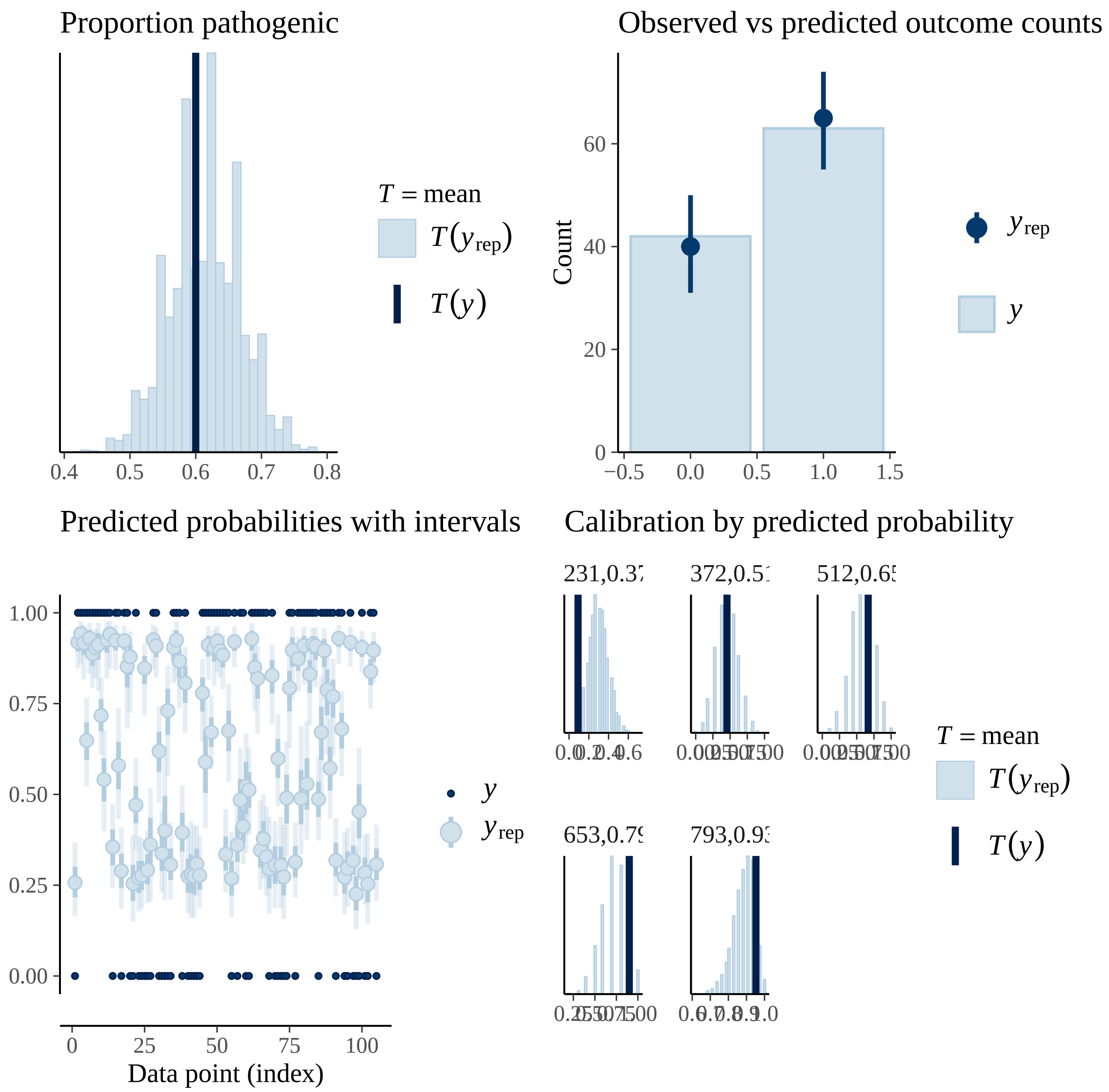


**Figure S4:** Posterior predictive checks for the distribution of the proportion of pathogenic variants (top left), the observed vs the predicted y=1 counts (top right), predicted probabilities of observed pathogenic variants (bottom left) and mean of predicted probabilities grouped by probability range (bottom right).

##### 3.1 Other Model Diagnostics

We performed twoo additional model diagnostics. First, we looked at the posterior distribution of the threshold parameter $U$ to see if the model identified a meaningful boundary between interface and non-interface regimes (Figure S5). Second, we examined the posterior probability assignments vs the various distance measures that we used (Figures S6 and S7).

While standard posterior predictive checks confirmed the model reproduces the observed outcome distribution (Figure SX), examination of the mixture components revealed that the model could not identify a meaningful interface boundary (Figure S5). The the posterior pathogenicity distribution is consistent with the interface being important (Figure S6 and S7), but is also consistent with this information being present in AM and EVE.

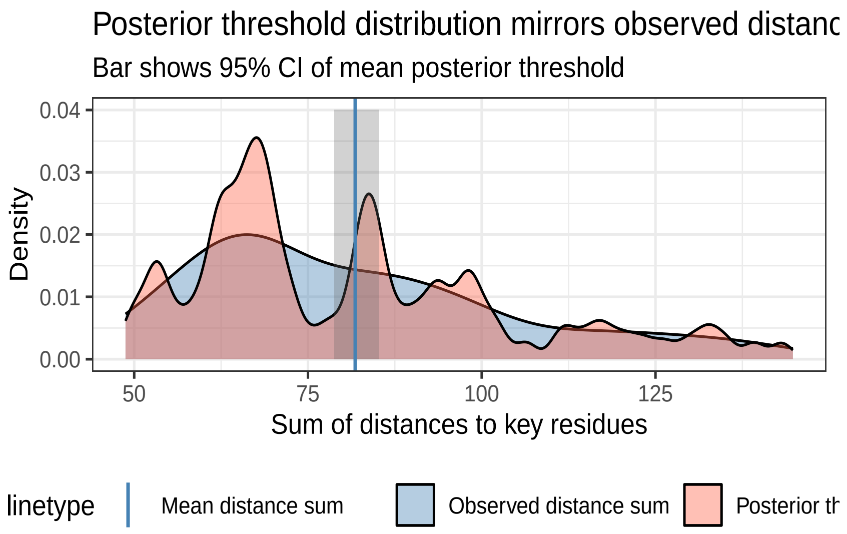


**Figure S5:** Distribution of the posterior distribution of the thresholds (the $U$ parameter) relative to the sums of the distances. The 95 credible interval (gray bar) of the mean posterior threshold overlaps with the mean of the observed distance sums (dashed line), and the overall distribution of posterior thresholds (tomato) closely follows the distribution of observed distance sums (steelblue). This shows that the model is not identifying a meaningful threshold that separates interface and non-interface regimes, but rather is reflecting the overall distribution of distances in the data.

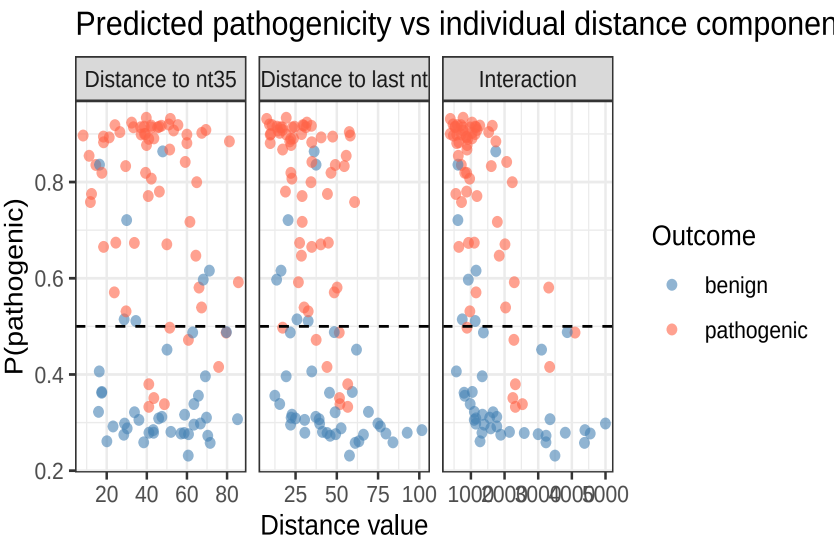


**Figure S6:** Posterior probability of pathogenicity as a function of the distance to nt35 (leftmost panel), last nucleotide (middle panel), and the interaction (rightmost panel). Points are colored by observed pathogenicity status (steelblue for benign, tomato for pathogenic). The predicted probability of pathogenicity shows a relationship to the last nucleotide and the interaction.

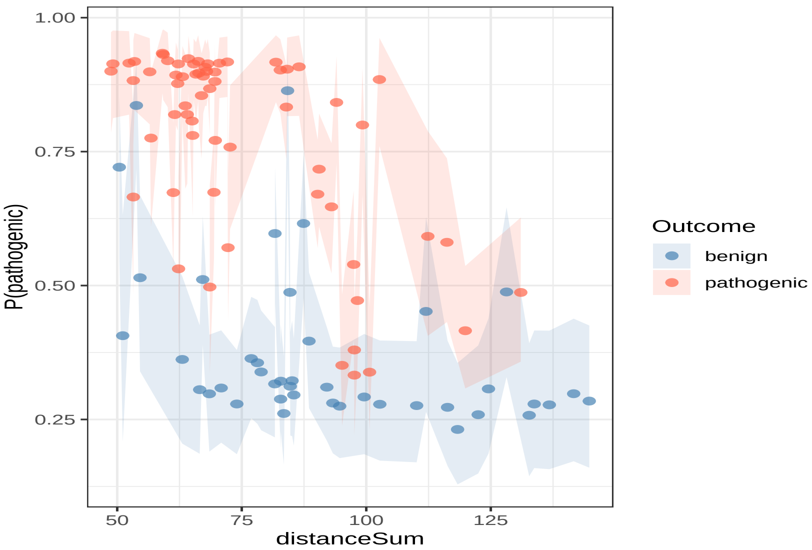


**Figure S7:** Posterior probability of pathogenicity as a function of the sums of distances to the two key residues. The ribbons indicate the posterior probability around each prediction.

### 4 Model Assessment

Once we have a working model and data, we need to check first how well we are able to fit relevant data with this model and to check that the model that we have really reflects the model that we are intending to fit.

We will address these two things simulating fake, yet realistic data. This is done by simulating data from the prior predictive distribution, fitting the data using the model, and using simulation based calibration [Talts et al., 2020] to evaluate the model.

Based on our priors, we generate observations:

$$\tilde{\theta} \sim p(\theta)$$
$$\tilde{y} \sim p(y|\tilde{\theta})$$

and then obtain $N$ posterior draws from the posterior distribution

$$\{\theta_1, ..., \theta_N\} \sim p(\theta|\tilde{y})$$

##### 4.1 Computational Faithfulness

To check the ability of our model to computational generated datasets, we will check the fitted datasets for the following metrics ([Vehtari et al., 2021b]):

- EBFMI (energy Bayesian fraction of missing information): this metric checks if the energy distribution of the Markov Chains is well behaved. If this metric is below 0.2, this suggests that the Markov Chains are not exploring the posterior well,
- Tree Depth: the Hamiltonian Monte Carlo method implemented in Stan uses a tree structure to explore the posterior. If the maximum tree depth is reached, this suggests that the posterior is difficult to explore and that we might need to increase the maximum tree depth or re-parameterize the model,
- no divergences: sometimes the posterior can have an exceedingly difficult geometry for the Markov Chains to explore, which can lead to biased estimates. Luckily, for the Hamiltonian Monte Carlo method implemented in Stan, these difficult geometries lead to divergences which are clearly identified and which we can use to diagnose and fix problems ([Betancourt, 2017]).

| Parameter | Percent |
|---|---|
| EBFMI | 0% |
| MaxTreeDepthReached | 0% |
| NumDivergences | 0% |

**Table S2:** Computational Faithfulness Metrics

##### 4.2 Simulation Based Calibration

The idea of simulation based calibration is based on the fact that the prior should have an identical distribution to the average of the model over the entire joint distribution:

$$p(\theta) = \int p(\theta, \tilde{y}, \tilde{\theta}) d\tilde{y} d\tilde{\theta} = \int p(\theta|\tilde{y}) p(\tilde{y}|\tilde{\theta}) p(\tilde{\theta}) d\tilde{y} d\tilde{\theta}$$

This means that any discrepancy between the prior and the averaged posterior over the data would suggest a problem in the model or the computation. We check the consistency between the prior and the data averaged posterior by using ranks. For each parameter we can define a rank function which would give us the number of posterior draws that are smaller than the simulated parameter:

$$r = \text{rank}(\tilde{\theta}, \{\theta_1, ..., \theta_N\}) = \sum_{n=1}^{N} \mathrm{I}(\theta_n < \tilde{\theta})$$

Talts et al [Talts et al., 2020] showed that $r$ is uniformly distributed on $[0, N]$, and we can use this to test the consistency of each parameter. We can obtain these ranks using simulated data, as we would have the true value as well as the posterior values. If the rank distribution systematically differ from uniformity for a parameter, this would suggest issues such as overdispersion, under-dispersion or bias.

For our case, as shown below there was no systematic deviation from the uniform distribution for any of the parameters (Figure S8).

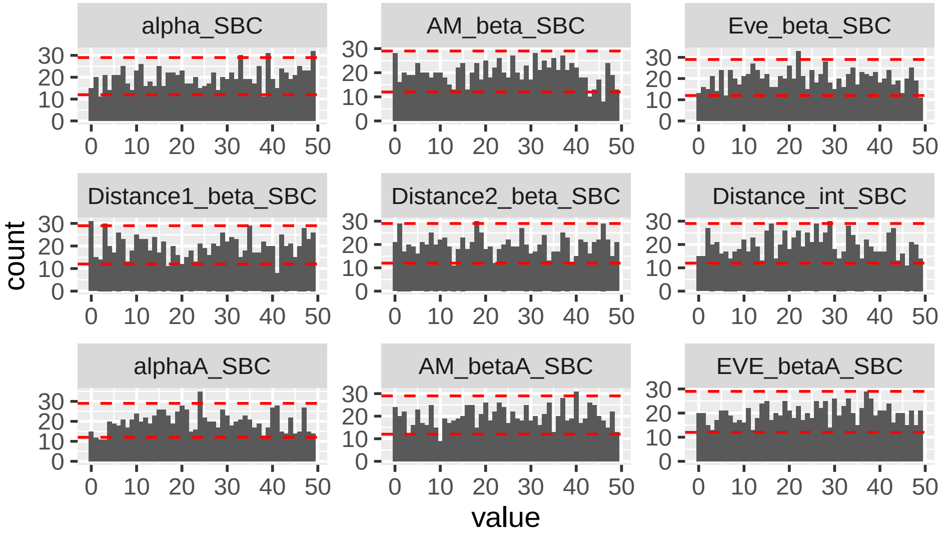


**Figure S8:** SBC.

#### 5 Model Sensitivity

To study the behavior of the posterior distribution in terms of bias and uncertainty, we will use two measurements. The first one is the *posterior z-score*:

$$z = \frac{\theta_{post} - \theta}{\sigma_{post}} \tag{S29}$$

where $\theta_{post}$ is the posterior mean for the parameter, $\theta$ the true value, and $\sigma_{post}$ the posterior standard deviation of the parameter. The difference between the true and the estimated value is scaled by the standard deviation, and hence this estimates how close the posterior estimate is to the true value. The further the $z$ is from zero, the further the posterior distribution is concentrated away from the true value.

The second measurement is called the *posterior contraction*:

$$c = 1 - \frac{\sigma^2_{post}}{\sigma^2_{prior}} \tag{S30}$$

and it shows how much the information from the likelihood, reflected in the posterior variance, $\sigma^2_{post}$, adds to the information from the prior, $\sigma^2_{prior}$. A $c$ value close to 1 indicates that the posterior variance is smaller than the prior variance and hence that the likelihood is very informative, whereas a $c$ value close to 0 suggests that the prior and posterior variances are close to each other and that the data has not added much information.

We can visualize the interplay between the behaviors of the *posterior z-score* and the *posterior contraction* with the following graph (Figure S9, modified from [Schad et al., 2019, Betancourt, 2020]):

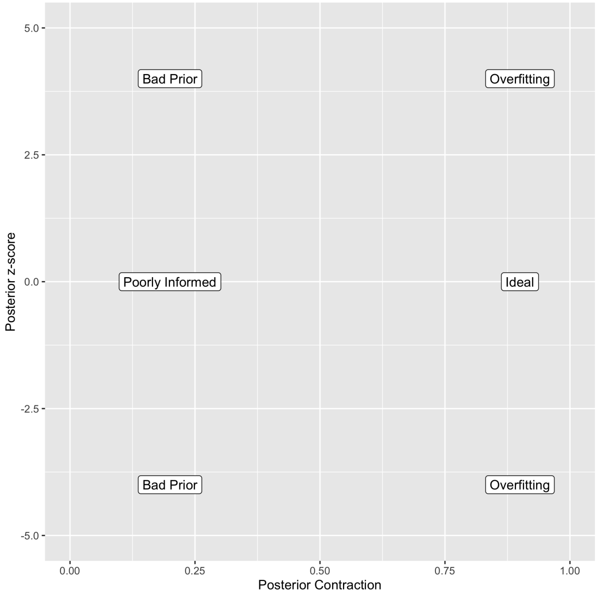


**Figure S9:** Interplay between posterior contraction and posterior-zscore. The labels indicate the situations one might encounter: high contraction indicates informative data, with a z-core close to zero indicating ideal behavior (informative data and accurate model) but large-zscore indicating over-fitting (informative data but biased inference); low contraction indicates less informative data, with a small z-score meaning that the observations do not inform the inference, and a large z-score indicating that the prior model is away from the true value.

The behavior for datasets with 105 observations is shown in Figure S10.

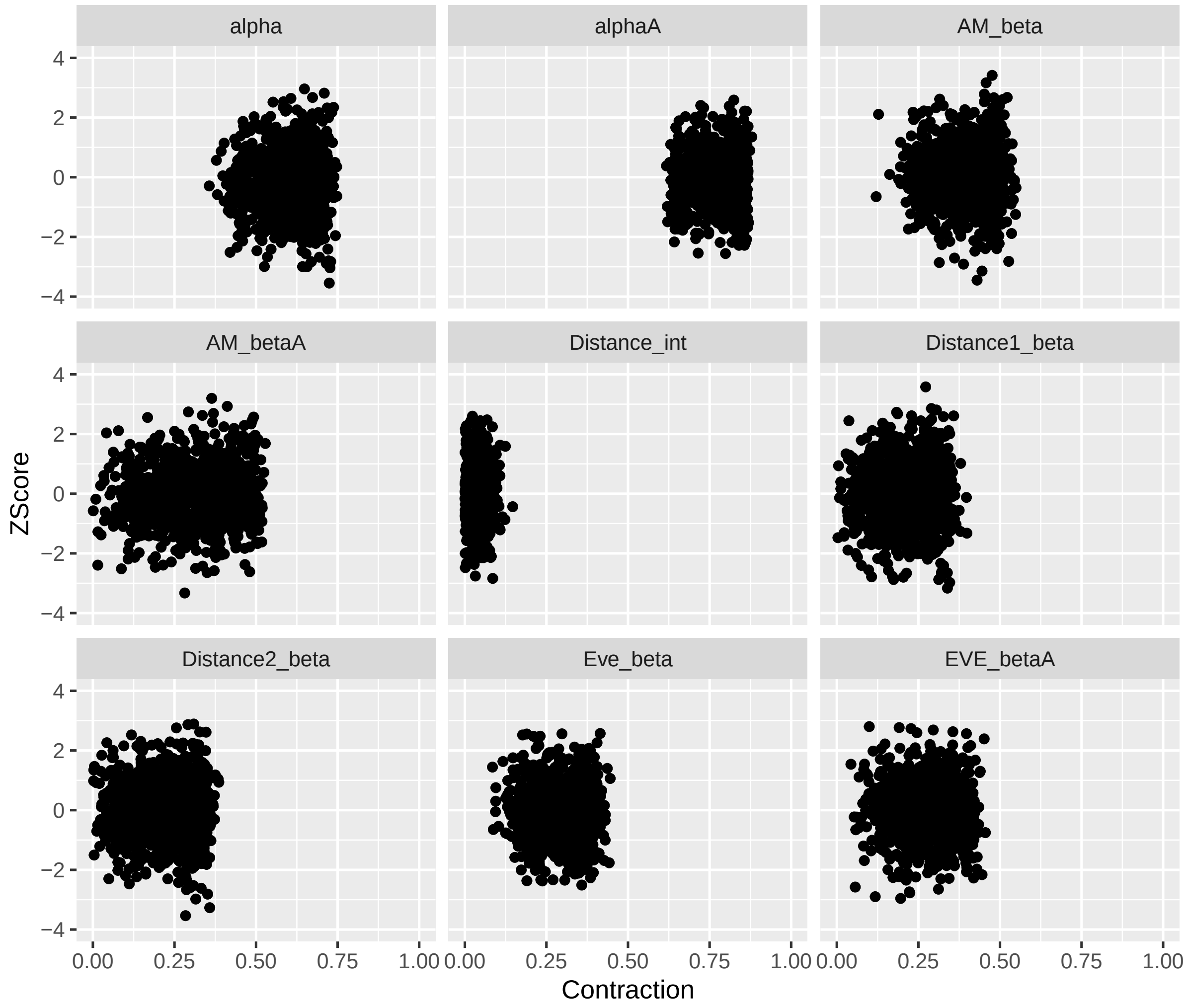


**Figure S10:** Posterior z-score vs posterior contraction for each parameter and each simulated dataset with 105 observations.

### 6 Comparison with Other Models

Both model estimate parameters for AM, EVE and the intercept, and have similar estimates, whereas the distance metrics for the AM+EVE+Distance models are not identified (Figure S11). The AM + EVE model outperformed the extended distance model on all three metrics — MSE (0.1142 vs 0.1152), ELPD LOO (-40.93 vs -44.88), and effective parameters (p LOO 1.47 vs 2.19), all of this suggests that the distance metrics add complexity but do not improve performance (Table **??**).

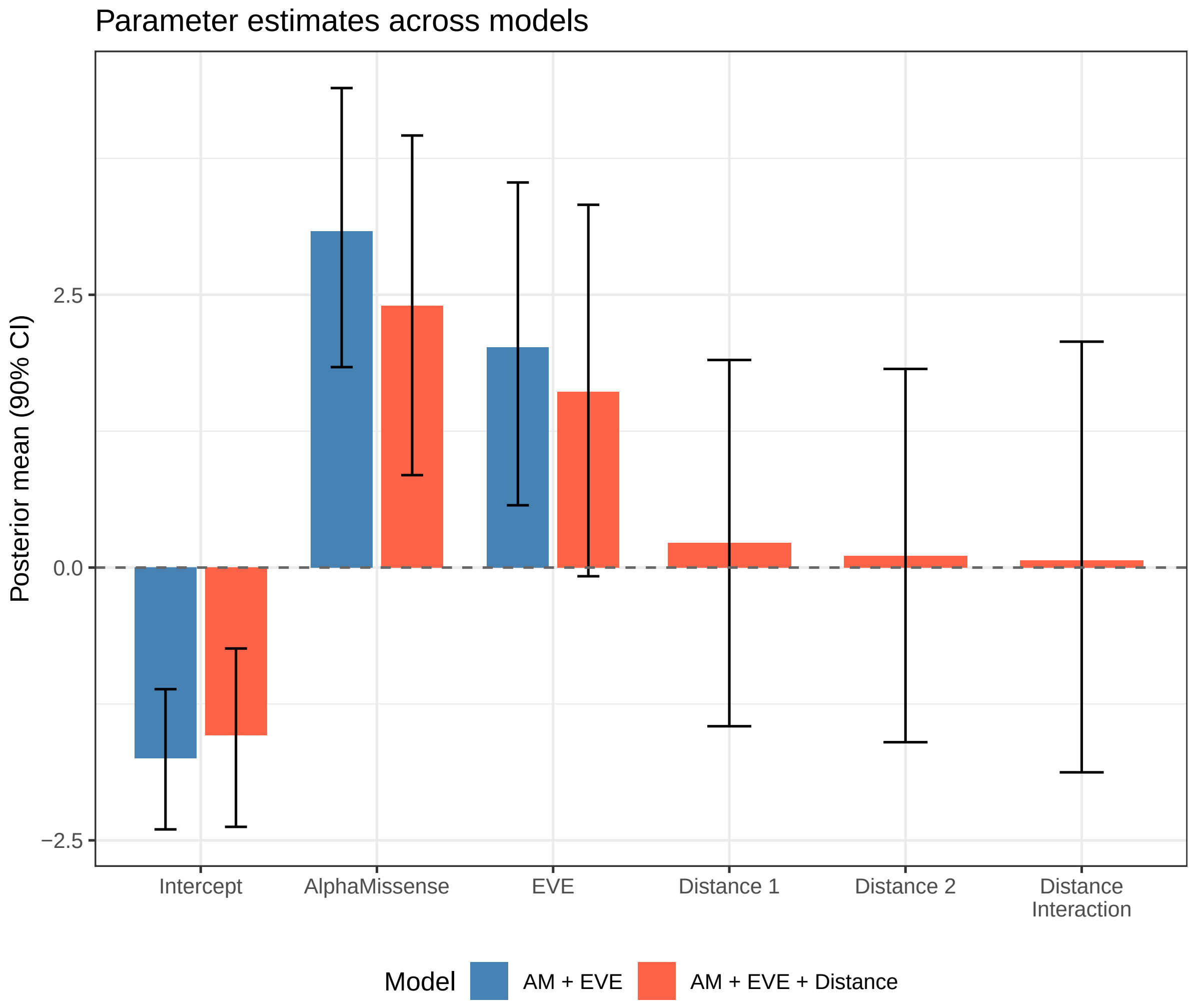


**Figure S11:** Posterior mean estimates and 90% credible intervals for the AM + EVE model and the AM + EVE + Distance model. Intercept, AlphaMissense, and EVE parameters are shared across both models. Distance 1, Distance 2, and Distance Interaction parameters are present only in the extended model. Dashed line indicates zero.

**Table S3:** Model comparison by mean squared error (MSE) and leave-one-out cross-validation (LOO). ELPD LOO is the expected log pointwise predictive density; higher values indicate better predictive performance. p LOO is the effective number of parameters.

| Model | MSE | ELPD LOO | SE | p LOO |
|---|---|---|---|---|
| AM | 0.1231 | -42.80 | 4.29 | 1.17 |
| EVE | 0.1426 | -48.36 | 3.89 | 1.17 |
| AM + EVE | 0.1140 | -40.78 | 4.72 | 1.39 |
| AM + EVE + Distance | 0.1150 | -44.80 | 4.06 | 2.14 |

### 7 Power Analysis

### 8 Description and Validation of Missing Data Imputation Model

For the missing data imputation we notice the following about the relationship between AlphaMissense and Eve:

They are correlated:

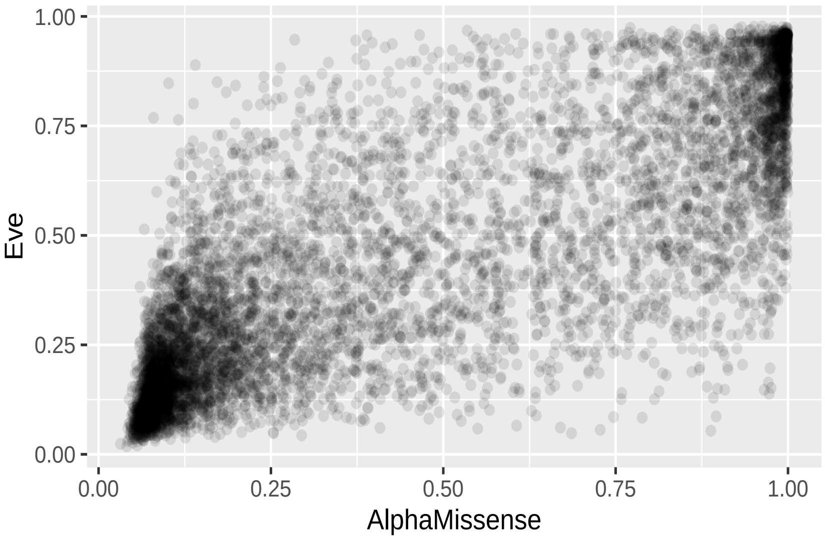


**Figure S12:** EVE values as a function of AM values.

With the mean of EVE increasing with increase in AlphaMissense, and the variance having a concave shape, with higher variance for intermediate values of AlphaMissense:

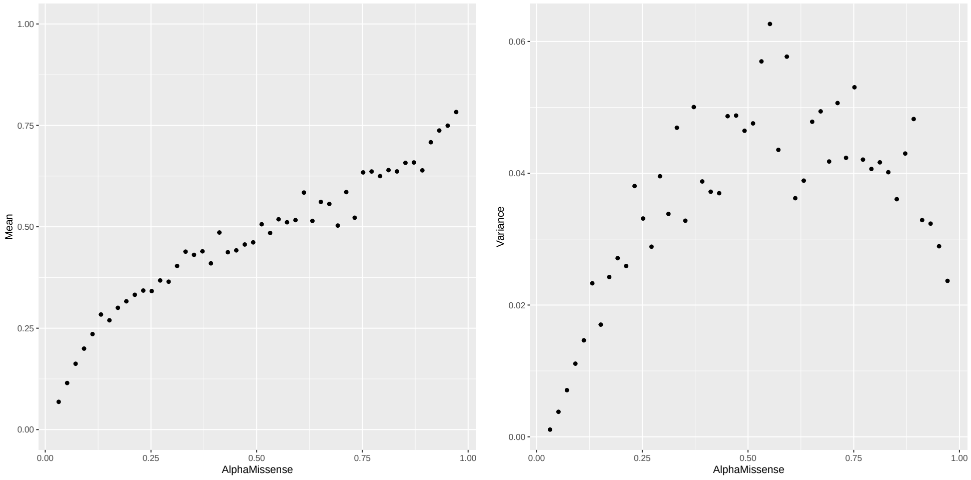


**Figure S13:** Mean of EVE values as a function of AM (left) and variance of EVE values as a function of AM (right).

#### 8.1 Model

Marginal distribution of AlphaMissense $AM$:

$$AM \sim \mathrm{Beta}(\alpha_2, \beta_2) \tag{S31}$$

Conditional distribution of EVE given AlphaMissense:

$$EVE \mid AM \sim \mathrm{Beta}(\alpha_1(AM), \beta_1(AM)) \tag{S32}$$

where we define the mean and the precision as follows:

$$\mu(AM) = \mathrm{logit}^{-1}(\mu_0 + \mu_1\, AM) \tag{S33}$$

$$\phi(AM) = \exp\Big(\phi_0 + \phi_1\, AM + \phi_2\, (AM - m)^2\Big) \tag{S34}$$

And use those as part of an alternative parameterization of the beta distribution,

$$\alpha_1(AM) = \mu(AM) \cdot \phi(AM) \tag{S35}$$

$$\beta_1(AM) = \big(1 - \mu(AM)\big) \cdot \phi(AM) \tag{S36}$$

We have $\mathbb{E}[EVE \mid AM] = \mu(AM) = \mathrm{logit}^{-1}(\mu_0 + \mu_1\, AM)$, is monotonically increasing when $\mu_1 > 0$, while staying in $[0, 1]$. The quadratic term $(AM - m)^2$ in $\log \phi$ allows the conditional variance to have a concave shape—higher in the middle $(m)$ of the $AM$ range and lower at the extremes.

For cases where the $EVE$ values are missing but $AM$ values are observed, we will impute the missing $EVE$ values using the above model jointly with the other parameters,

$$AM_i \sim \mathrm{Beta}(\alpha_2, \beta_2) \tag{S37}$$

$$EVE_i^{\mathrm{imp}} \sim \mathrm{Beta}\big(\alpha_1(AM_i), \beta_1(AM_i)\big) \tag{S38}$$

**Priors**

$$\alpha_2 \sim \text{Beta}(1, 1) \tag{S39}$$

$$\beta_2 \sim \text{Beta}(1, 1) \tag{S40}$$

$$\mu_0 \sim \mathcal{N}(-2, 1) \tag{S41}$$

$$\mu_1 \sim \text{LogNormal}(1, 0.5) \tag{S42}$$

$$\phi_0 \sim \mathcal{N}(1, 0.5) \tag{S43}$$

$$\phi_1 \sim \mathcal{N}(1, 0.5) \tag{S44}$$

$$\phi_2 \sim \text{LogNormal}(0, 1) \tag{S45}$$

$$m \sim \mathcal{N}(0.5, 0.05) \tag{S46}$$

###### 8.1.1 Prior Predictive Checks

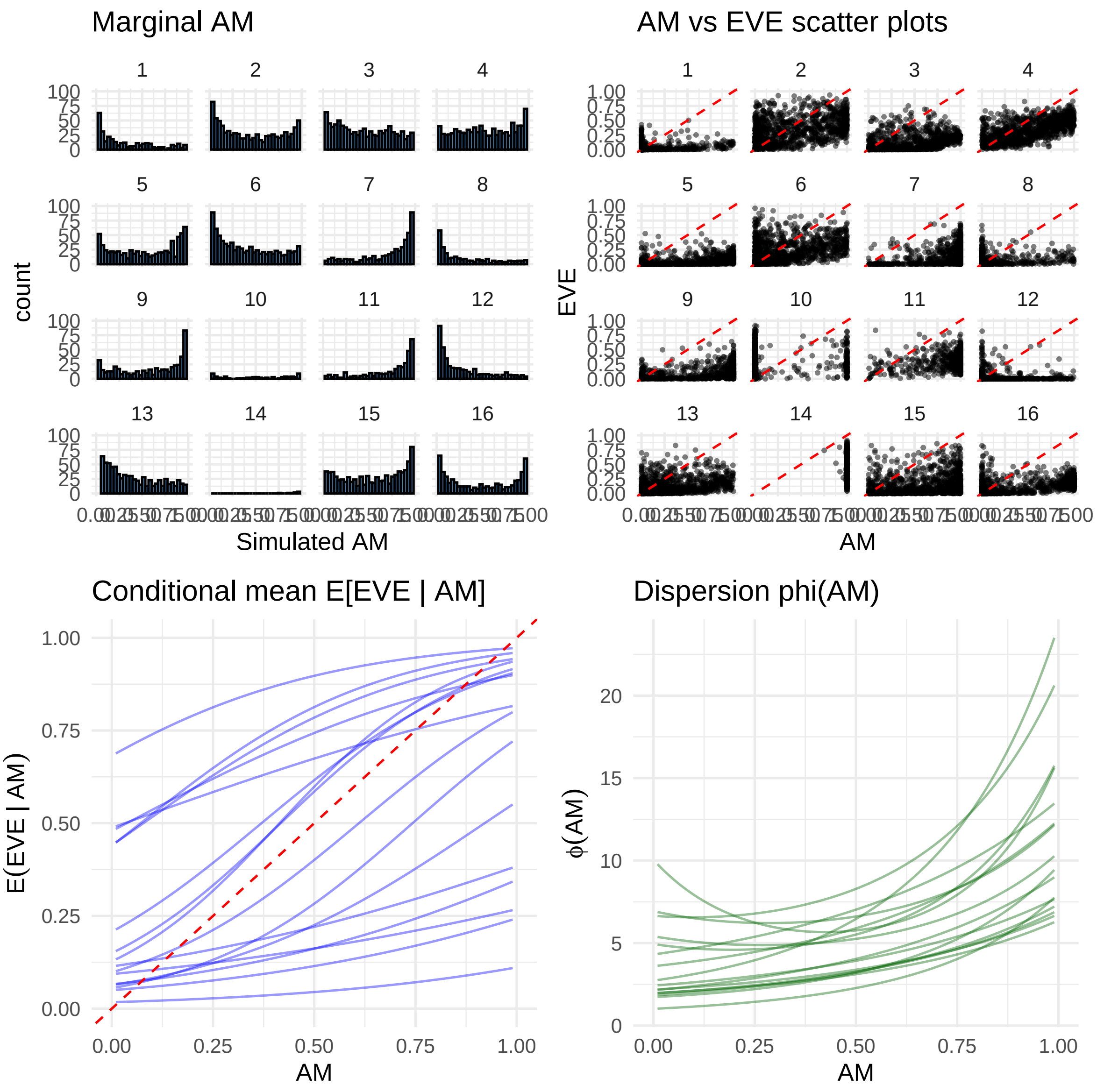


**Figure S14:** Prior predictive checks: histograms of simulated AM, should have U shaped; scatter plots of simulated AM vs EVE; conditional mean increases as AM increases; variance should be lower in the two ends

#### 8.2 Posterior Predictive Checks

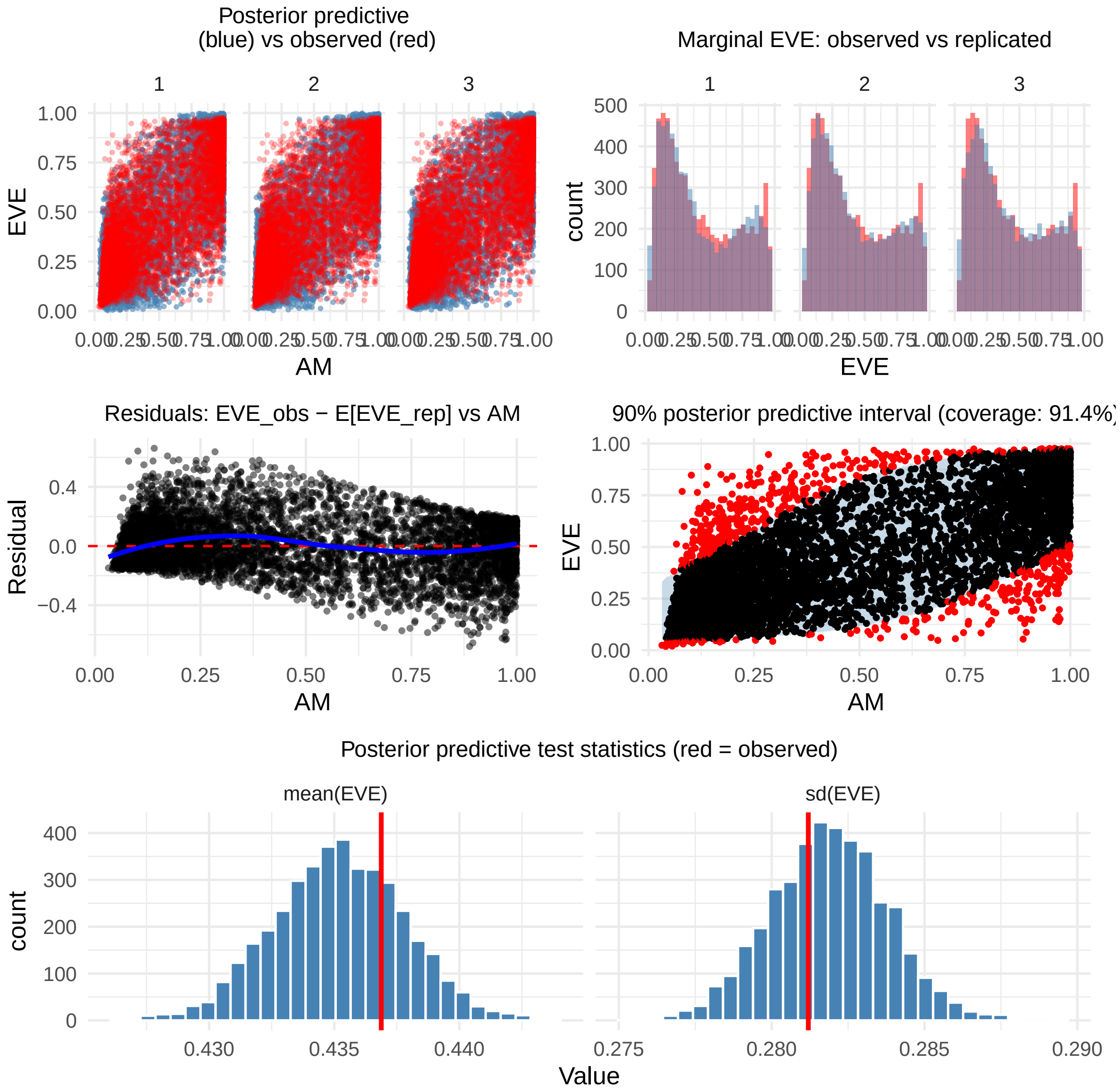


**Figure S15:** Posterior predictive checks: (top left) scatter plot of observed AM vs replicated EVE, should look similar to observed AM vs observed EVE; (top right) histogram of replicated EVE vs histogram of observed EVE, should look similar; (middle left) residuals of observed EVE - posterior mean EVE vs AM, should be centered around 0 with no clear pattern, just shows a very small change; (middle right) 90% posterior predictive interval for each data point, with points colored by whether the observed value is inside the interval, should have around 90% coverage; (bottom) test statistics (mean of EVE, sd of EVE) for replicated data vs observed value, observed value should be in the bulk of the replicated distribution. They all are except the correlation.

#### 8.3 Final Imputed Observations

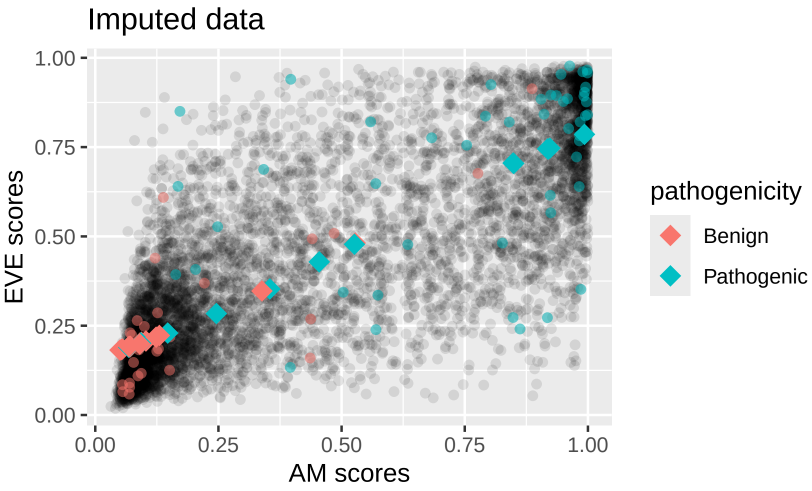


**Figure S16:** Imputed EVE data points (in diamonds)

**Table S4:** Imputed EVE scores

| | AM_pathogenicity_scores | EVE_inferred |
|---|---|---|
| DARS_H_280_L | 0.9193 | 0.7442810 |
| GARS_E_738_K | 0.0936 | 0.2036816 |
| GARS_L_32_P | 0.0502 | 0.1817727 |
| GARS_P_42_A | 0.0529 | 0.1831204 |
| GARS_V_91_I | 0.0699 | 0.1916016 |
| HARS_Q_14_K | 0.5276 | 0.4814361 |
| KARS_T_595_S | 0.0714 | 0.1920660 |
| LARS_E_1137_K | 0.1028 | 0.2071582 |
| LARS_K_1155_E | 0.0906 | 0.2012064 |
| LARS_L_37_V | 0.0551 | 0.1846272 |
| NARS_K_60_T | 0.5261 | 0.4780658 |
| NARS_R_238_Q | 0.9182 | 0.7451865 |
| NARS_R_322_L | 0.9930 | 0.7857354 |
| NARS_R_329_Q | 0.9228 | 0.7469482 |
| NARS_R_545_C | 0.8481 | 0.7053226 |
| NARS_T_17_M | 0.0926 | 0.2018887 |
| NARS_V_226_L | 0.4546 | 0.4286610 |
| QARS_A_77_T | 0.0701 | 0.1902790 |
| QARS_G_45_V | 0.3542 | 0.3528481 |
| QARS_M_1_T | 0.2458 | 0.2840092 |
| QARS_Y_57_H | 0.8495 | 0.7042290 |
| RARS_D_2_G | 0.0927 | 0.2014409 |
| RARS_M_1_T | 0.1470 | 0.2294286 |
| RARS_M_1_V | 0.0679 | 0.1896925 |
| RARS_R_28_W | 0.1235 | 0.2178403 |
| TARS_E_77_D | 0.1301 | 0.2226107 |
| TARS_G_21_D | 0.0905 | 0.1985692 |
| VARS_R_181_C | 0.3383 | 0.3472044 |
| WARS_L_9_V | 0.0685 | 0.1919735 |

### 9 Session Information

We used R v. 4.5.3 [R Core Team, 2026] and the following R packages: bayesplot v. 1.15.0 [Gabry et al., 2019, Gabry and Mahr, 2025], cmdstanr v. 0.9.0 [Gabry et al., 2025], doParallel v. 1.0.17 [Corporation and Weston, 2022], dplyr v. 1.2.1 [Wickham et al., 2026], foreach v. 1.5.2 [Microsoft and Weston, 2022], ggplot2 v. 4.0.3 [Wickham, 2016], ggpubr v. 0.6.3 [Kassambara, 2026], iterators v. 1.0.14 [Analytics and Weston, 2022], kableExtra v. 1.4.0 [Zhu, 2024], knitr v. 1.51 [Xie, 2014, Xie, 2015, Xie, 2025], LaplacesDemon v. 16.1.8 [Statisticat and LLC., 2026[illegible] Statisticat and LLC., 2026a, Statisticat and LLC., 2026c, Statisticat and LLC., 2026d], loo v. 2.9.0 [Vehtari et al., 2017, Yao et al., 2018, Magnusson et al., 2019a, Magnusson et al., 2019b, Paananen et al., 2021, Vehtari et al., 2024a, Vehtari et al., 2025, Sivula et al., 2025], MASS v. 7.3.65 [Venables and Ripley, 2002], patchwork v. 1.3.2 [Pedersen, 2025], posterior v. 1.7.0 [Vehtari et al., 2021a, Lambert and Vehtari, 2022, Margossian et al., 2024, Vehtari et al., 2024b, Bürkner et al., 2026], reshape2 v. 1.4.5 [Wickham, 2007], scales v. 1.4.0 [Wickham et al., 2025a], tidyr v. 1.3.2 [Wickham et al., 2025b], tidytable v. 0.11.2 [Fairbanks, 2024], xtable v. 1.8.8

[Dahl et al., 2026].
