## Supporting Information Tables for "Structural distance at the tRNA synthetase active site interface predicts pathogenicity but is captured by AlphaMissense and EVE except among score-ambiguous variants"

**S1 Table.** Comparison of data sources. “Unique Proteins” shows how many synthetases are included. “Total Mutations” for UniProt includes variants downloaded from UniProt that were not included in the final analysis due to either a lack of pathogenicity annotation or conflicting annotations; see S2 Table for more detail. “Average Pathogenicity Score” is shown for EVE and Alpha Missense (AM), but not for UniProt or Distance because they do not have quantitative values assigned for pathogenicity of mutations.

| Data | EVE | AM | UniProt | Distance |
|---|---|---|---|---|
| Unique Proteins | 14 | 19 | 19 | 13 |
| Total Mutations | 203460 | 283157 | 16941 | 8440 |
| Average Pathogenicity Score | 0.541 | 0.599 | | |
| Median Pathogenicity Score | 0.552 | 0.661 | | |

**S2 Table.** Mutation data summary. Individual proteins and corresponding variant counts for which structural distance information was available are included. The number of mutations with an Alpha Missense (AM) and EVE score are listed, but EVE lacks data for NARS. UniProt annotations (pathogenic and benign) are also included. “With UniProt annotation” includes any variant with an annotation on UniProt under “clinical significance.” Some annotations contained conflicting interpretations of pathogenicity, so “Pathogenic (UniProt)” and “Benign (UniProt)” show the variants with consistent annotations on UniProt; these were the only variants included in the “true” pathogenicity classifications. See methods for how annotations were classified as pathogenic or benign.

| Protein name | UniProt ID | Total mutations | With AM score | With EVE score | With UniProt annotation | Pathogenic (UniProt) | Benign (UniProt) |
|---|---|---|---|---|---|---|---|
| CARS | P49589 | 730 | 730 | 700 | 9 | 4 | 1 |
| DARS | P14868 | 529 | 529 | 424 | 24 | 7 | 0 |
| GARS | P41250 | 1045 | 1045 | 828 | 245 | 6 | 8 |
| HARS | P12081 | 655 | 655 | 519 | 161 | 2 | 6 |
| KARS | Q15046 | 800 | 800 | 560 | 32 | 4 | 2 |
| LARS | Q9P2J5 | 1124 | 1124 | 973 | 21 | 3 | 4 |
| NARS | O43776 | 577 | 577 | 0 | 19 | 7 | 0 |
| QARS | P47897 | 1194 | 1194 | 909 | 276 | 5 | 4 |
| RARS | P54136 | 721 | 721 | 567 | 27 | 5 | 3 |
| TARS | P26639 | 740 | 740 | 640 | 9 | 2 | 4 |
| VARS | P26640 | 1224 | 1224 | 958 | 54 | 12 | 2 |

| | | | | | | | |
|---|---|---|---|---|---|---|---|
| WARS | P23381 | 408 | 408 | 311 | 7 | 2 | 1 |
| YARS | P54577 | 675 | 675 | 660 | 151 | 4 | 7 |
| All Proteins | | 10422 | 10422 | 8049 | 1035 | 63 | 42 |

**S3 Table.** Summary of Sequence Alignment Quality. Comparing aligned distance data sequences to UniProt data sequences. "Total Rows" indicates the number of variants available in the dataset for each protein (this includes only variants that exist both in UniProt/Alpha Missense and in the pdb sequence used for distance calculations); these can include multiple variants in the same position with different substitutions. "Available AA Pairs" represents the number of amino acids paired for each sequence alignment (this includes only one amino acid per position). "Matching AAs" shows the number of amino acid pairs in each sequence alignment that were identical. "Percent Matching" shows the ratio of matching amino acids to total amino acid pairs available.

| Protein | Total Rows | Available AA Pairs | Matching AAs | Percent Matching |
|---|---|---|---|---|
| CARS | 730 | 481 | 396 | 82.3 |
| DARS | 529 | 322 | 253 | 78.6 |
| GARS | 1045 | 509 | 440 | 86.4 |
| HARS | 655 | 322 | 97 | 30.1 |
| KARS | 800 | 456 | 418 | 91.7 |
| LARS | 1124 | 748 | 621 | 83 |
| NARS | 577 | 368 | 317 | 86.1 |
| QARS | 1194 | 572 | 513 | 89.7 |
| RARS | 721 | 447 | 374 | 83.7 |
| TARS | 740 | 468 | 387 | 82.7 |
| VARS | 1224 | 776 | 617 | 79.5 |
| WARS | 408 | 274 | 206 | 75.2 |
| YARS | 675 | 335 | 284 | 84.8 |
